# Connecting Thiamine Availability to the Microbial Community Composition in Chinook Salmon Spawning Habitats of the Sacramento River Basin

**DOI:** 10.1101/2023.08.22.554313

**Authors:** Christopher P. Suffridge, Kelly C. Shannon, H. Matthews, R. Johnson, C. Jeffres, N. Mantua, Abigail E. Ward, E. Holmes, J. Kindopp, M. Aidoo, F. Colwell

## Abstract

Thiamine Deficiency Complex (TDC) is a major emerging threat to global populations of culturally and economically important populations of salmonids. Salmonid eggs and embryos can assimilate exogenous thiamine, and evidence suggests that microbial communities in benthic environments can produce substantial amounts of thiamine. We therefore hypothesize that microbially produced thiamine in both riverine surface water and hyporheic zones could serve to rescue early life stages of salmonids suffering from TDC. The distributions of thiamine and its metabolically related compounds (dTRCs) have never been determined in freshwater systems. Similarly, the microbial cycling of these compounds has never been investigated. Here we determine that all dTRCs are present in femto-picomolar concentrations across diverse salmon spawning habitats in California’s Sacramento River system. We observed that thiamine concentrations in the Sacramento River are orders of magnitude lower than marine environments, indicating substantial differences in thiamine cycling between these two environments. Our data suggest that the hyporheic zone is likely the source of thiamine to the overlying surface water. Temporal variations in dTRC concentration were observed where highest concentrations were seen when Chinook salmon were actively spawning. Significant correlations were identified between the richness of differentially abundant ASVs and dTRC concentrations. The influence of these ASVs on dTRC concentrations provide evidence of dTRC cycling by microbes in the hyporheic zone, which would influence the conditions where embryonic salmon incubate. Together, these results indicate a connection between microbial communities in freshwater habitats and the availability of thiamine to spawning TDC-impacted California Central Valley Chinook salmon.

## IMPORTANCE

Pacific salmon are keystone species with considerable economic importance and immeasurable cultural significance to Pacific Northwest indigenous peoples. Thiamine Deficiency Complex has recently been diagnosed as an emerging threat to the health and stability of multiple populations of salmonids ranging from California to Alaska. Microbial biosynthesis is the major source of thiamine in marine and aquatic environments. Despite this importance, the concentrations of thiamine and the identities of the microbial communities that cycle it are largely unknown. Here we investigate microbial communities and their relationship to thiamine in Chinook salmon spawning habitats in California’s Sacramento River system to gain an understanding of how thiamine availability impacts salmonids suffering from Thiamine Deficiency Complex.

## INTRODUCTION

Thiamine Deficiency Complex (TDC) is an emerging threat to global populations of salmonid fish (1–3). TDC is caused by chronic low systemic concentrations of thiamine (vitamin B1), resulting in early life stage mortality (1, 4). The occurrence of thiamine deficiency has been globally documented in habitats including The Baltic Sea, the New York Finger Lakes, the Laurentian Great Lakes, the Yukon River in Alaska, the Sacramento-San Joaquin River watershed in California, and coastal rivers in Oregon (4–7). Many studies have linked TDC occurrence to salmonid consumption of planktivorous fish species containing the thiamolytic enzyme, thiaminase (1, 5, 8–15). Clupeid prey with observed thiaminase activity include anchovies, herring, sprat, and alewife, which can each contribute to the bulk of salmonid diets in various aquatic ecosystems. Consumption of these clupeids have been linked to thiamine deficiency (1, 6, 8, 12). A second, yet not mutually-exclusive, hypothesis links TDC incidence to the consumption of prey fish with high lipid and energy densities, such as sprat, which causes oxidative stress and resulting thiamine depletion in predators (16). Incidences of TDC have increased globally in multitudes of animal species (2) and it is now considered a major emerging threat to the stability of many globally distributed populations of salmonids and their associated fisheries (1, 2).

TDC has recently been diagnosed for the first time in Chinook salmon populations spawning in the California Central Valley, including those spawning in the Sacramento River system (Figure 1)(6). Symptoms of TDC in this system were first observed in recently hatched fry in hatcheries in 2020 (6). These juvenile fish displayed common symptoms of TDC including lethargy, abnormal swimming patterns, loss of equilibrium, and death (6). Fisheries managers were able to successfully mitigate TDC in the hatchery populations by treating fry and eggs with thiamine baths and thiamine injections (6, 17). In this incidence, TDC was hypothesized to be linked to an unprecedented dominance of Northern Anchovy in the diets of these salmon (6).

**Figure 1.**
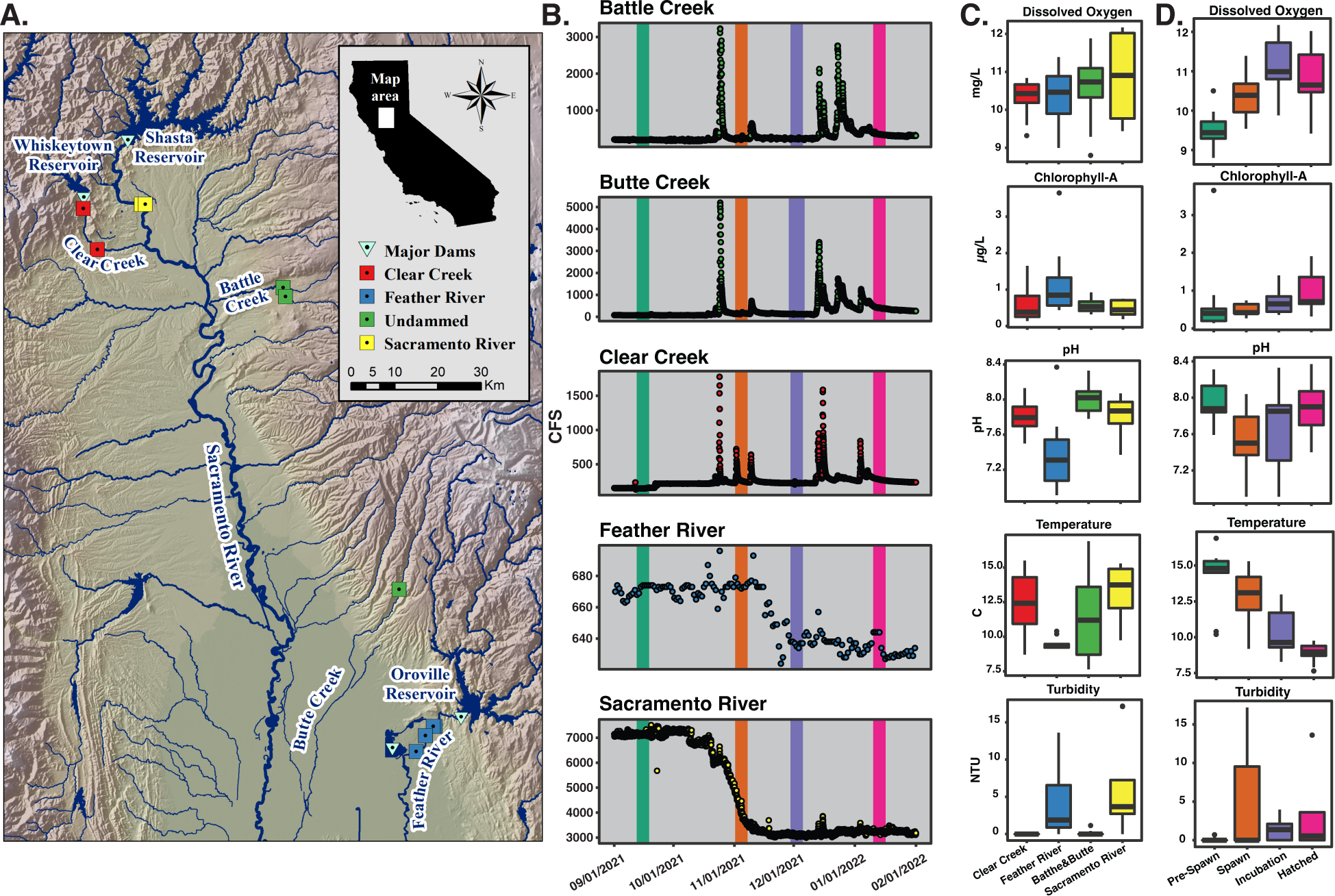
Geographic, physical, and chemical parameters of sampling sites. A. Sample locations in the Sacramento River watershed. Sample locations are coded by color throughout the manuscript: Sacramento River (yellow), Clear Creek (red), Feather River (blue), and Battle and Butte Creeks (green). B. Hydrological data from each stream from September 2021 to February 2022 (cubic feet per second). Colored bars represent sampling times, and these colors are used throughout the manuscript: Pre-Spawn (teal), Spawn (orange), Incubation (purple), and Hatched (pink). C. Box plots of water chemistry data binned by river. D. Box plots of water chemistry data binned by sampling event.

Anchovy populations in the California Current ecosystem historically undergo boom and bust cycles, often alternating in dominance with sardines, presumably in response to varying ocean conditions (18, 19). We theorize that microbial thiamine production in Chinook salmon spawning habitats may naturally mitigate the impacts of ocean sourced TDC.

While post-embryonic salmonids primarily acquire thiamine through dietary intake, all life stages of salmonids (including eggs and embryos) can acquire thiamine directly from the dissolved pool in the surrounding environment via uptake over the gills and diffusion, respectively (1, 7, 20). The degree to which salmon could be supplemented by dissolved thiamine at each life stage is unknown. It is unlikely that direct thiamine uptake from the dissolved pool represents a significant source of this vitamin once salmonids are actively feeding (regardless of developmental stage or habitat occupied) because their prey likely satisfies the majority of their metabolic thiamine demands. However, during adult salmon’s freshwater migration and spawning life stages when adults are fasting, and for eggs and embryos incubating in riverine sediments, direct uptake of exogenous thiamine may represent a major, critically important thiamine source. Past research has established that thiamine deficiency in salmonid embryos is maternally transmitted (21). Therefore, the environmental dissolved thiamine pool could mitigate the pathological effects of TDC experienced by spawning female salmonids.

Vitamin B1 is an essential coenzyme universally required by all domains of life, yet it is mainly biosynthesized by a relatively small subset of microorganisms and most microbes are auxotrophs (obligately require) for the vitamin (22, 23). Vitamin B1 functions as a cofactor for roughly 2% of all characterized enzymes that require a cofactor, and is essential for both catabolic and anabolic central carbon metabolism and branched-chain amino acid synthesis (24). Thiamine is a heterocyclic molecule containing pyrimidine and thiazole moieties which are synthesized by two separate biosynthesis pathways and ligated to form thiamine, which is phosphorylated through multiple steps to its active state, thiamine pyrophosphate (25). Multiple metabolically relevant thiamine related compounds (TRCs) must be simultaneously analyzed to fully assess environmental microbial thiamine cycling (26). The generalized microbial thiamine cycle is visually outlined in Figure 2; the concentration of each compound can be considered as a proxy for different metabolic processes in the thiamine cycle. The primary biosynthetic precursors to thiamine (referred to as B1 hereafter) are the pyrimidine compound 4-amino-5-hydroxymethyl-2-methylpyrimidine (HMP) and the thiazole compound 5-(2-hydroxyethyl)-4-methyl-1,3-thiazole-2-carboxylic acid (cHET) (25, 27). These compounds are solely produced through biosynthetic processes and can be considered proxies for active thiamine biosynthesis. It has been reported that the thiazole compound 4-methyl-5-thiazoleethanol (HET) is both a B1 biosynthetic precursor and degradation product (produced by both intracellular and extracellular processes) that can be utilized through a salvage pathway to complete cellular B1 biosynthesis (22, 25, 27–32). Similarly, cells are known to be able to utilize exogenous sources of the B1 pyrimidine degradation product 4-amino-5-aminomethyl-2-methylpyrimidine (AmMP) through a salvage pathway to complete B1 biosynthesis, and thus AmMP can be viewed as a proxy for thiamine degradation (32).

**Figure 2.**
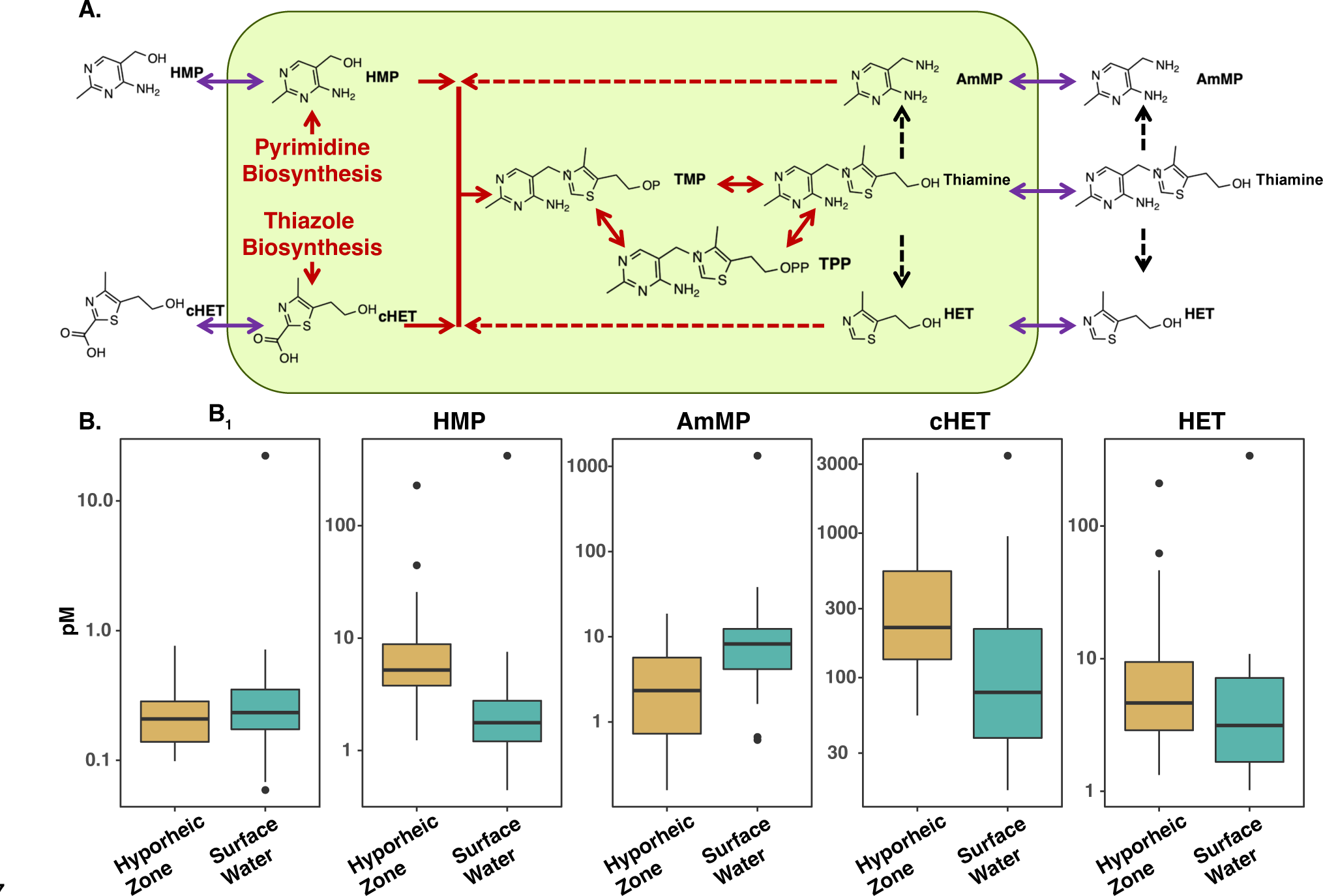
dTRCs within the Sacramento River watershed. A. Summary of microbial thiamine cycle pathways. The green rectangle represents a generic microbial cell. Solid red arrows represent cellular biosynthetic processes. Dashed red arrows represent cellular salvage pathways. Dashed black arrows represent degradation processes. Purple double headed arrows represent cellular excretion and uptake processes. In some cases, arrows represent multiple enzymatic and/or abiotic processes. B Distribution of dTRCs within the Sacramento River watershed. Boxplots compare concentrations (picomolar; pM) of all river SW (blue) and HZ (tan) samples of each dTRC.

The concentrations of dissolved TRCs (dTRCs) in freshwater environments are largely unknown. Less than 50 discrete freshwater dissolved B1 measurements exist worldwide, and concentrations are reported in the picomolar range (33–35). Measurements of B1 from Lake Tahoe, California and the Peconic River Estuary in New York range between 12-190 pM however these studies used bioassay and HPLC based methods which cannot accurately discriminate between all TRCs and are therefore likely overestimates (33, 34). Tovar-Sanchez and colleagues (2016) used a LCMS based approach to measure B1 in the Moulouya River, Africa and observed a mean concentration of 4.5 pM (35). The concentrations of other dTRCs have never been determined in freshwater systems.

The distribution of dTRCs have been more extensively studied in marine environments ranging from oligotrophic gyres to eutrophic coastal environments (26, 36–38). In these environments concentrations of each dTRC are within the picomolar range (26). Organisms that harbor thiamine biosynthesis genes span the domains of life and include bacteria, archaea, eukaryotic algae, plants, and fungi (37, 39). However, species within each domain differ in thiamine biosynthetic capacity. Some contain a complete repertoire of biosynthesis genes and produce thiamine *de novo* (prototrophs), whereas others must rely on salvage pathways to complete thiamine biosynthesis through cellular transport of exogenous dissolved thiamine precursor compounds or intact thiamine (auxotrophs) (22, 40). In fact, both culture based and genome based research provides evidence that the majority of marine microorganisms are thiamine auxotrophs (22, 27–29, 32, 37, 41). Specifically, many of these organisms lack the capability to synthesize one, or both, of thiamine’s precursor moieties (e.g., HMP, cHET) and therefore must acquire these compounds (or analogous degradation products. E.g., AmMP, HET) from the exogenous dissolved pool to complete thiamine biosynthesis (22, 27–29, 32, 41). The relative abundance of thiamine prototrophic and auxotrophic algae and bacterioplankton can influence dTRC concentrations in the marine water column (26). The rates of dTRC uptake and production are largely unknown, but given the widespread nature of thiamine auxotrophy, microbial demand for dTRCs is likely substantial. Microbial dTRC production balances removal. Both culture- and genome-based studies in marine systems have provided evidence that many species of free-living bacterioplankton, including cyanobacteria, produce dissolved thiamine (37, 42). Further, the Peconic River was shown to be a source of thiamine for the coastal ecosystem of Long Island, NY, displaying a potential connection between thiamine cycling in freshwater, estuary, and marine systems (34). Sediment microbial communities also produce substantial amounts of thiamine and release it to the exogenous dissolved pool. For example, one study in the Santa Monica Basin offshore of Los Angeles, California estimated that marine sediments could produce thiamine at 0.7 nmol•m^-2^•day^-1^ (43), and in freshwater systems research has shown correlative relationships between concentrations of sediment-sourced trace metals and thiamine, suggesting that the microbial inhabitants of river sediments produce thiamine to the overlying water (35). Therefore, the observed standing stock concentrations of dTRCs in any environment represent an equilibrium between thiamine production and removal.

Here we report the first investigation of the distribution of dTRCs coupled with microbial community structure in freshwater systems, with the objective of determining the availability of thiamine to Chinook salmon impacted by TDC. The relationship between microbial community compositions and concentrations of dTRCs was assessed in both the surface water (SW) and hyporheic zone (HZ) of the Sacramento River basin, a freshwater, lotic environment where adult Chinook salmon spawn and their offspring rear. We hypothesize that (1) microbial community compositions of the SW and HZ of the Sacramento River and its tributaries impacts the concentrations of dTRCs and (2) both dTRC concentrations and compositions of microbial communities of each sample type (HZ and SW) differ by region of the basin and time points related to the spawning of Sacramento River fall-run Chinook salmon. We seek to gain an understanding of how environmental changes, including natural and anthropogenic, can impact the structure of aquatic microbial communities and thus the availability of thiamine to higher trophic levels. Further, we assess which taxonomic annotations (microbial taxa) correlate most significantly with dTRC concentrations to help infer how the relative abundance of certain microbial taxa can impact dTRC cycling. Additionally, we aim to provide a baseline mechanistic understanding of how microbially-produced TRCs in aquatic environments could serve as natural mitigation for both adult and embryonic Chinook salmon impacted by TDC. Pacific salmon are keystone species (44) with considerable economic importance and immeasurable cultural significance to Pacific Northwest indigenous peoples (45, 46). Therefore, examining the structure of microbial communities that produce and consume TRCs in rivers where Chinook salmon spawn is tremendously important.

## METHODS

### Sample collection

Samples were collected in September, November, and December of 2021 and in January of 2022. These dates were chosen to bracket key events for fall-run Sacramento River Chinook salmon, and they correspond to before spawning (Pre-Spawn), during active spawning (Spawn), during the salmon egg incubation (Incubation), and after the eggs had hatched but the embryonic fish had not yet left the gravel and started feeding exogenously (Hatched)(47). Eleven sites within the Sacramento River watershed were sampled at each of the four time points (Figure 1). Sites (listed North to South) were located on the Sacramento River below Shasta and Keswick dams (SAC, SSC), Clear Creek below the Whiskeytown Dam (CCM, CCH), North Fork Battle Creek (NFB), South Fork Battle Creek (SFB), Butte Creek (BCC), and the Feather River below the Oroville Dam (FRU, FRD; FR2, FR3 only at the December sampling). Sites were chosen to capture a gradient of anthropogenic influence. Clear Creek, the Sacramento River, and the Feather River sites are all downstream of major high head dams and reservoirs that fully prevent upstream fish passage. North and South forks of Battle Creek and Butte Creek are impacted by small dams and water diversions and lack major reservoirs. Chinook salmon are known to actively spawn at all sites. The sites on the Feather River in Oroville are adjacent to a hatchery. FRU is upstream of the hatchery while sites FRD, FR2, and FR3 are downstream.

Samples were collected from both the river SW and HZ at each site. SW samples were collected directly from a depth of 0.5 m using 1 L amber HDPE bottles (Nalgene). HZ samples were collected by driving a standpipe 30 cm into the unconsolidated river gravels in areas typical of salmon spawning. The bottom 20 cm of standpipe was perforated to allow sample collection from the HZ. A teflon tube was then inserted into the standpipe, and the sample was extracted using a hand powered pump. Care was taken to extract all overlying river water from the standpipe prior to sample collection. Samples were collected into 1 L amber HDPE bottles. Collected samples from both the SW and HZ were then prefiltered using a 100 μm mesh to remove all metazoans and large suspended particles. Sampling occurred in daylight hours over the course of one day for each sampling event. All sample collection equipment was cleaned as appropriate for trace organic compound sampling which has been described previously (48).

Briefly, all bottles, tubing, and pumps were acid washed using 1 M hydrochloric acid, followed by rinses with MilliQ water and finally methanol to remove all organics. Sampling equipment was thoroughly rinsed with sample before sample collection. Water samples were stored on ice until filtration occurred.

Concurrent with water sampling a YSI EXO2 Multiparameter Water Quality Sonde was used to collect river metadata. Data was collected for temperature (TEMP; °C), dissolved oxygen percent saturation (DO; %), dissolved oxygen concentration (mg•L^-1^), turbidity (Turb; NTU), total dissolved solids (TDS), chlorophyll-a concentration (CHL; μg•L^-1^), relative blue-green algae concentration (BGA; μg•L^-1^), electrical conductivity (EC; μS•cm^-1^) and specific conductance (i.e., temperature-standardized conductivity, SPC; μS•cm^-1^), salinity (Sal; PSU), and pH. Data for river discharge (CFS) was downloaded from the California Department of Water Resources California Data Exchange Center (https://cdec.water.ca.gov/dynamicapp/wsSensorData).

River SW and HZ water samples were processed for dTRC and microbial community analysis as has been described previously (26, 48). Briefly, gentle peristaltic filtration (30 ml•min^-1^) was used to collect all cells and particles on a 0.22 μm Sterivex filter (PES membrane, Millipore, Burlington, MA, USA). Microbial biomass on the Steviex filter was used for 16S rRNA gene amplicon microbial community diversity analysis. After filtration, the filter was blown dry, capped, and stored at -80 °C until analysis. One liter of cell free filtrate was collected in an acid washed, methanol rinsed, amber HDPE bottle for dTRC analysis. The dTRCs in the filtrate were stabilized with the addition of 1 ml of 1 M hydrochloric acid, and the filtrate was stored at -20 °C until analysis. Samples were processed using a portable field laboratory to allow for processing to occur the same day as collection. All samples were transported frozen to Oregon State University for analysis.

### dTRC analysis

dTRCs were extracted from river and HZ water as previously described (26, 48). Samples were thawed and adjusted to pH 6.5 using 1 M HCl or 1 M NaOH. dTRCs were extracted from the river and HZ water matrix using a solid phase extraction with Bondesil C_18_ resin (Agilent). Samples were passed over 8 ml of resin at the rate of 1 ml•min^−1^. Previous research has determined that all dTRCs were retained on the resin (26). Compounds were eluted from the column using 12 ml of LCMS grade methanol, which was then concentrated by evaporation to 250 μl using a blow-down nitrogen drier (Glass Col). This provides a six order of magnitude concentration factor between the environmental sample concentration and the concentration analyzed on the LCMS. Hydrophobic organic compounds that were co-extracted by the solid phase extraction were removed using a liquid phase extraction with 1:1 volume of chloroform. Samples were then stored at −80 °C until LCMS analysis.

A liquid chromatography mass spectrometry (LCMS) method was utilized to simultaneously measure dTRC concentrations (26). Analysis was conducted using an Applied Biosystems 4000 Q-Trap triple quadrupole mass spectrometer with an ESI interface coupled to a Shimadzu LC-20AD liquid chromatograph. Applied Biosystems Analyst and ABSciex Multiquant software were used for instrument operation and sample quantification. A Poroshell 120 PFP, 3 × 150 mm 2.7 μm HPLC column (Agilent) with a Poroshell 120 PFP, 2 × 5mm, 2.7 μm guard column (Agilent) was used for chromatographic separations. The column temperature was isocratic at 40 °C. HPLC mobile phases were MS grade water (Fisher) with 0.1% formic acid and MS grade acetonitrile (Fisher) with 0.1% formic acid. A 15-min binary gradient was used with a flow rate of 200 μl•min^−1^ and an initial concentration of 3% acetonitrile ramping to 100% acetonitrile in 7 min and column re-equilibration at 3% acetonitrile for 6 min. A third HPLC pump with a flow rate of 100 μl•min^−1^ acetonitrile (0.1% formic acid) was connected to a mixing tee post column to increase the ionization efficiency as most dTRCs elute from the column in the aqueous phase of the gradient. The ESI source used a spray voltage of 5,200 V and a source temperature of 450 °C. Curtain gas pressure was set at 30 PSI. The mass spectrometer was run in positive ion mode. Compound specific information including MRM parameters, column retention times, and limits of detection have been previously published (26). The sample injection volume was 10 μl, and samples were analyzed in triplicate. Samples were indiscriminately randomized prior to analysis. To compensate for matrix effects, ^13^C-labeled thiamin was used as an internal standard. LCMS analysis was conducted at the Oregon State University Mass Spectrometry Core Facility.

### DNA extraction, amplification, and sequencing preparation

Microbial DNA was extracted from Sterivex filters representing approximately 1 L of SW or HZ water per sample site. Using an autoclave-sterilized scalpel, each Sterivex filter membrane was sectioned in half (representing approximately 500 ml of sample) and placed in Qiagen DNeasy Powersoil DNA isolation kit bead-beating tubes (Qiagen; Hilden, Germany) for immediate DNA extraction and the other half for storage at -80 °C. Sterivex filters in each bead-beating tube were dissolved by adding 0.1 ml of phenol chloroform in a fume hood and the remainder of the DNA extraction protocol was performed following the manufacturer’s SOP. All Sterivex extraction steps prior to phenol chloroform addition (done in chemical fume hood) were carried out in a laminar flow hood. Negative controls were performed in parallel by adding 0.25 ml of PCR-grade water to bead-beating tubes and carrying out all DNA extraction and PCR steps.

The V4-V5 hypervariable region of the bacterial and archaeal 16S SSU rRNA gene was PCR-amplified with indexed 515F-806R 16S primers (49, 50). Two separate high throughput sequencing runs were performed on the Feather River sample sites and the remainder of the sample sites. PCR for Feather River samples was performed in 25 μl double-well reactions containing 200 nM of forward and reverse primers, 12.5 μl of Platinum II Hot-Start PCR Master Mix (2X) (Invitrogen Waltham, Massachusetts), 1 μl of undiluted DNA template, and PCR-grade sterile water to a total volume of 25 μl. The HZ FRU Spawn time period sample was PCR-amplified with 6 μl template due to a low extraction yield and HZ Incubation-time period FR3 sample was PCR-amplified with 1 μl 1:5-diluted sample for proper PCR amplification. Volumes of all other samples contained 1 μl of undiluted template DNA. Thermocycler conditions were performed following the PCR master mix manufacturer’s procedure and consisted of the following steps for all samples: 2 min initial denaturation at 94 °C, 30-35 cycles of denaturation at 94 °C for 15 s, annealing at 60 °C for 15 s, and extension at 72 °C, followed 10 min at 72 °C for a final non-cycled hold. Feather River samples were pooled to an equimolar concentration and sequenced at the Oregon State University Center for Quantitative Life Sciences (CQLS) for 2×250 bp Illumina MiSeq high throughput sequencing in June, 2022 and the remainder of samples were pooled to an equimolar concentration and sequenced at the CQLS using the same sequencing platform in October, 2022.

### Bioinformatics and statistics of microbial community data

The CQLS demultiplexed all raw MiSeq reads and trimmed the majority of adaptor sequences. Final Illumina adaptor trimming and initial quality filtering was performed with Trim Galore (v0.6.7) to drop reads with Phred scores of <20. FastQC (v0.11.9) was used to generate read quality reports and MultiQC (v1.14; (51)) was used for visualization of combined reports. All further computational work was performed in RStudio (v4.1.1; (52)). DADA2 (v124.0; (53)) was used to perform final read filtering, contig creation, and read assignment of prokaryotic ASVs, following default parameters, with the exception of conducting pseudo-pooling at the denoising step to allow for better recognition of rarer ASVs in each sample based on prior information (54). The Silva (v138.1; (55)) 16S reference database was used to assign taxonomy to bacteria and archaea and the Phytoref (56) reference database was used to assign plastid taxonomy to ASVs that were identified as Chloroplast sequences by Silva alignment in DADA2. Decontam (v1.16.0; (57)) was used with default parameters to remove contaminant ASVs based on the existence of these ASVs in negative control samples.

A Phyloseq object was generated from the ASV count and taxonomy tables of bacteria, archaea, and Phytoref-assigned algae from DADA2 and by integrating an associated metadata table using the phyloseq package (v1.42.0; (58)). This Phyloseq object was then rarefied to an even depth of 8,087 reads, representing 17,733 total ASVs, and all samples were maintained. Separate Phyloseq objects were generated for total bacteria and archaea (excluding algae-assigned ASVs), HZ bacteria, archaea, and Phytoref-assigned algae, and SW bacteria, archaea, and Phytoref-assigned algae. Phyloseq was used to generate Bray-Curtis dissimilarity matrices for each Phyloseq object and vegan (v2.6-4; (59)) was used to find significant differences between groups (PERMANOVA), significant differences between group variances, and to generate vectors for NMDS plots with the “adonis2”, “betadisper” and “permutest”, and “gg_envfit” commands, respectively. All Bray-Curtis dissimilarity NMDS plots were visualized with ggplot2 (v3.4.0; (60)). Constrained analysis of principal coordinates (CAP) was performed with the “ordinate” command from Phyloseq on bacterial and archaeal communites (not algal) with the variables, region, time point, and sample type, and was plotted with the “plot_ordination” command from Phyloseq. The package MicrobiotaProcess (v1.8.2; (61)) was used to generate alpha diversity richness and Shannon diversity index (ASV richness and evenness) values of whole communities (bacteria, archaea, and eukaryotic algae) for boxplots with the command “get_alphaindex”. Kruskal Wallis tests on differences in Shannon diversity indices and on differences in dTRC concentrations between groups was performed with the command, “kruskal.test” from the stats (52) package.

Mantel tests were performed with the “mantel” command from vegan between transformed (Tukey’s ladder of powers), continuous dTRC concentration values and Bray-Curtis dissimilarity, calculated with the “distance” command from Phyloseq. Mantel tests were also performed between Bray-Curtis dissimilarity and Haversine distance by transforming latitude and longitude values of each sample site to Haversine distance values with the “distm” command from the package, geosphere (62)). Mantel tests were performed on SW and HZ bacterial, archaeal, and algal communities separately. Differential abundance was performed with ANCOM-BC (63)) with unrarefied phyloseq objects of hyporheic and SW samples separately, and with raw and continuous dTRC concentrations. The top ten most significant (lowest p-values; all < 0.05) ASVs in each log-fold change direction, associated with each dTRC, were used for further analysis. Correlations analyses were done with metadata transformed with Tukey’s ladder of powers, to maximize linearity between variables (64) and normalized to an ordinal scale with the equation: [(“#$%& − min(“#$%&))/(max(“#$%&) − min(“#$%&))] for Spearman correlations, where “value” is each observation of the variable of interest and min and max are the minimum and maximum values of that variable, respectively. Spearman correlation coefficients were calculated with the function, “cor”, p-values with the function, “cor.mtest”, and correlograms plotted with “corrplot”, all with the corrplot package (65)). Shannon diversity index values for the correlations analysis were produced by the “diversity” function from the microbiome package (66)) and raw richness measurements were taken by summing all ASV counts of differentially abundant ASVs from rarefied (see above) ASV count tables. dTRC concentration values were binned into “low”, “normal”, and “high” based on the 25th, 50th, and 75th percentile, respectively, of each dTRC to assess how the relative abundance of ASVs changed by dTRC status. ASV relative abundance values by metadata grouping variables were visualized and identified with the Phinch2 desktop application (v2.0.1; (67)).

## RESULTS

### Physical characteristics of the river environment

Samples were collected from eleven stations on five rivers within the Sacramento River watershed at four time points designed to bracket fall-run Chinook salmon spawning (Figure 1). Sample locations had a range of anthropogenic impacts. Heavily impacted sites were located on the Feather River, Clear Creek, and the Sacramento River. Sites on Battle Creek and Butte Creek were relatively less impacted, however upstream water diversion structures and other channel modifications do exist on these rivers. One measure of the degree of anthropogenic control of each tributary was demonstrated by the observed river flow rates which showed natural rain-event driven pulses in Battle, Butte, and Clear Creeks (Figure 1). Major rain events occurred immediately before the Spawn (November) and Hatched (January) sampling events (Figure 1).

In contrast, elevated flow pulses were not seen in the Feather and Sacramento River sites where the flow was regulated by the upstream high head dams in a bimodal nature with high flow (670 and 7000 CFS, respectively) before November and low flow (640 and 3000 CFS, respectively) after November. The rain events between November and January caused increases in turbidity at the Spawn and Hatched time points that were especially evident downstream of Lake Oroville and Lake Shasta in the Feather River and Sacramento River, respectively where turbidity reached 13.6 and 17.2 NTU, respectively (Figure 1).

Temporal and regional patterns were observed in the water chemistry. Temperature across all stations dropped from a high median of 14.8 °C in September to 8.9 °C in January (Figure 1). The Feather River was consistently the coldest and had the lowest variability in temperature of the tributaries with an observed temperature range of 9.2-10.4 °C (Figure 1). Conversely, the Battle and Butte Creek sites had the largest variability of observed temperatures with a range of 7.6-16.9 °C. Chlorophyll-a concentration was significantly negatively correlated with temperature, with concentrations across all stations increasing from a median of 0.4 μg•L^-1^ in September to 0.71 μg•L^-1^ in January (Figures 1,4). The Feather River sites had the highest observed chlorophyll-a concentrations ranging from 0.44-3.65 μg•L^-1^ (Figure 1). Across all sites median pH decreased from 7.88 in September to 7.5 in November, and then to 7.85 in January (Figure 1). The Feather River had the lowest pH range across all time points (excluding outliers) running from 6.91-7.54, while all other tributaries had similar pH ranges across all time points ranging from 7.37-8.33 (Figure 1). Dissolved oxygen concentrations increased from a median value of 9.44 mg•L^-1^ in September to 10.99 mg•L^-^1 in December before dropping to 10.65 mg•L^-1^ in January. No regional differences in dissolved oxygen were observed when samples were binned by tributary (Figure 1).

### Distribution of dTRCs

dTRCs were detected in the femto-picomolar range in both the HZ and the SW of the Sacramento River watershed. Samples from all stations and time points were binned based on sample type (HZ and SW) to understand the basin-scale relationships between thiamine cycling in the HZ and SW (Figure 2, Table 1). The B1 concentrations in both the HZ and SW were not significantly different (tests for significant differences between HZ and SW means done with paired t-tests; median values are 0.21 and 0.23 pM, respectively), and the interquartile ranges (IQR) of concentrations in the HZ and SW reflected this similarity with overlapping values of 0.14-0.29 pM and 0.17-0.35 pM (Figure 2, Table 1). The concentrations of thiamine’s biosynthetic precursors, HMP and cHET, did not vary significantly but were higher in the HZ than in the SW (Figure 2). The IQR of HMP in the HZ was 3.78-8.84 pM whereas the SW IQR was 1.21-2.77 pM, and median HMP concentration was 3 times higher in the HZ than the SW. Similarly, the median concentration of cHET was 2.8 times higher in the HZ than the SW. The IQRs for cHET in the HZ and SW were 133.67-546.33 pM and 38.22-219 pM, respectively (Figure 2, Table 1). Opposingly from the findings of HMP and cHET, the median concentration of AmMP, a thiamine degradation product, was 3.5 times higher in the SW than the HZ and the differences in means were significant (p-value<0.05; 8.25 and 2.33 pM, respectively). The AmMP IQR was 4.17-12.38 pM in the SW and 0.73-5.70 in the HZ. The IQRs of HET, both a thiamine degradation product and biosynthesis precursor, overlapped in the HZ and SW with values of 2.89-9.45 pM and 1.74-7.78 pM, respectively. The median of HET in the HZ was 4.64 pM and 3.14 pM in the SW. HET was frequently undetectable in the SW, and was only detected in the Feather River and the South Fork of Battle Creek (Figure S1, Table S1).

**Table 1.** Summary of HZ and SW dTRC picomolar concentrations. Data from this table is graphically presented in Figure 2. The complete dTRC dataset is presented in Table S1.

|  | B1 |  | HMP |  | AmMP |  | cHET |  | HET |  |
| --- | --- | --- | --- | --- | --- | --- | --- | --- | --- | --- |
|  | HZ | SW | HZ | SW | HZ | SW | HZ | SW | HZ | SW |
| **Minimum** | 0.10 | 0.06 | 1.24 | 0.45 | 0.16 | 0.61 | 54.73 | 16.67 | 1.33 | 1.02 |
| **Lower Quartile** | 0.14 | 0.17 | 3.78 | 1.21 | 0.73 | 4.17 | 133.67 | 38.22 | 2.89 | 1.74 |
| **Median** | 0.21 | 0.23 | 5.20 | 1.77 | 2.33 | 8.25 | 222.50 | 79.53 | 4.64 | 3.14 |
| **Upper Quartile** | 0.29 | 0.35 | 8.84 | 2.77 | 5.70 | 12.38 | 546.33 | 219.67 | 9.45 | 7.78 |
| **Maximum** | 0.76 | 22.30 | 227.67 | 418.33 | 18.63 | 1333.33 | 2623.33 | 3436.67 | 209.67 | 338.33 |

Multiple statistical outliers (Tukey Test) were identified at both the high and low concentration extremes in the HZ and SW pools for all dTRCs (Figure 2, Table 1). These outliers likely represent biologically significant events as has been previously established (26, 38). Further evidence for the biological relevance of these statistical outliers is that when concentrations are assessed on an individual station basis, South Fork Battle Creek was the source of the high extreme outliers in both the HZ and the SW at two separate sampling events: Spawn and Incubation (Figure S1, Table S1).

We observed a substantial temporal pattern in dTRC concentrations in both the HZ and SW across all samples where the concentrations at Spawn were higher than the preceding and following sample points for all dTRCs but cHET (Figure 3). Temporal variability in dTRC concentrations was assessed by binning samples from all stations by sample time point (Figure 3). We determined that time point bins, rather than regional bins, were able to resolve a greater amount of the observed variation in dTRC concentrations as Kruskal-Wallis tests only show significant regional variability between regions for B1 in the SW, whereas this test found significant temporal variability for HMP in the SW, AmMP in the HZ and SW, and cHET in the HZ and SW (p-values<0.05). Additionally we were able to qualitatively validate these results by visually comparing boxplot medians and IQRs which showed no clear regional trends, but a clear, repeatable temporal trend (Figures 3, S1).

**Figure 3.**
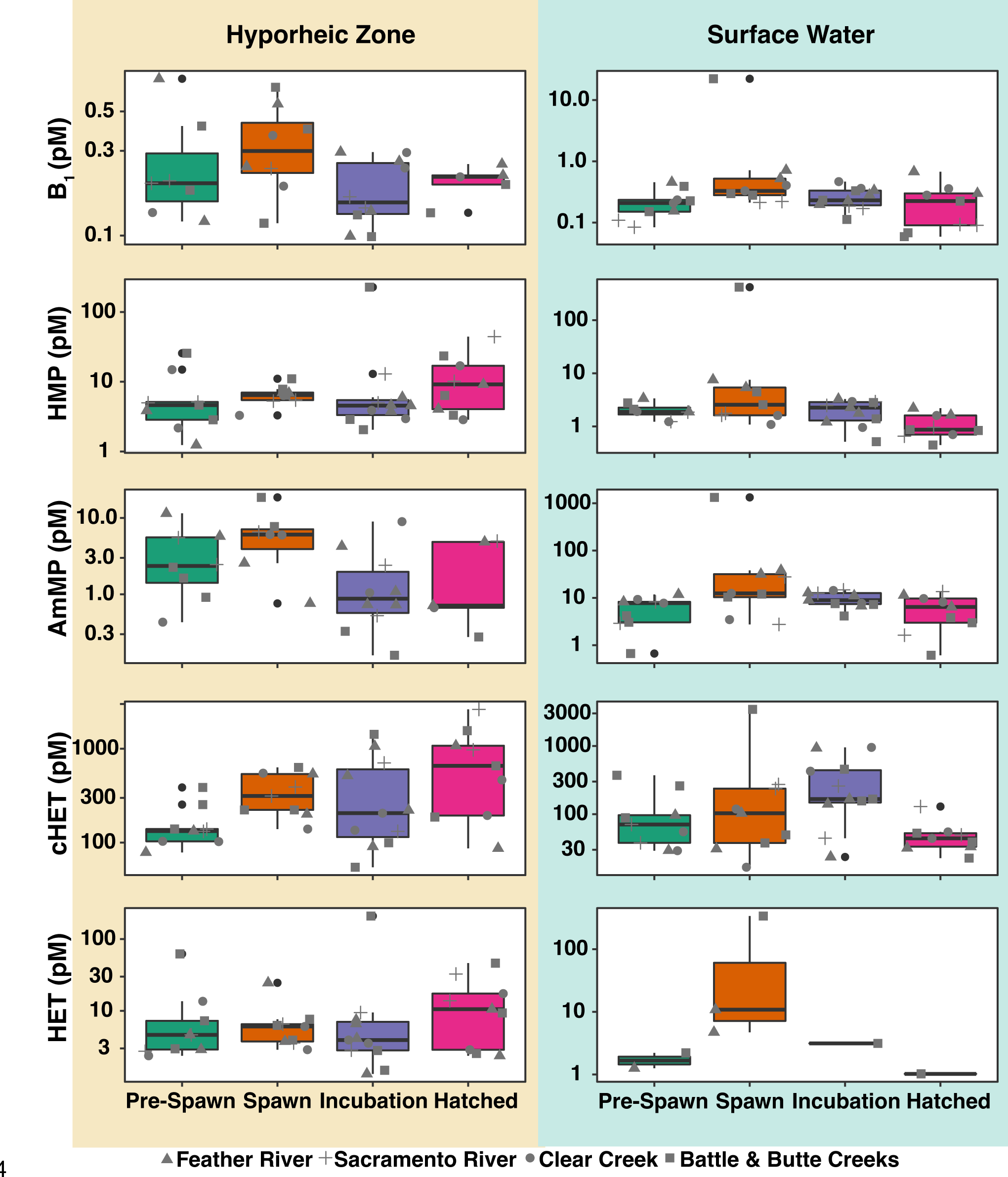
Temporal variations in dTRC concentrations. Box plots of Hyporheic Zone (tan) Surface Water (blue) dTRC concentrations are binned by time point. Box colors represent sampling times. Individual points are overplotted and shapes indicate tributary: Feather River (triangle), Sacramento River (plus), Clear Creek (circle), and Battle and Butte Creeks (square).

The temporal trend we observed in the HZ showed an increase in median dTRC concentration from Pre-Spawn (0.2 pM B1, 4.63 pM HMP, 2.36 pM AmMP, 133.0 pM cHET, and 4.63 pM HET) to Spawn (0.31 pM B1, 6.43 pM HMP, 6.06 pM AmMP, 314.0 pM cHET, and 6.06 pM HET) followed by a decrease of median dTRC concentration at the Incubation time point (0.15 pM B1, 4.54 pM HMP, 0.89 pM AmMP, 206.0 pM cHET, and 3.93pM HET) (Figure 3). The median concentration of all dTRCs except AmMP in the HZ at the Hatched time point increased from the Incubation time point (0.21 pM B1, 9.18 pM HMP, 0.71 pM AmMP, 657.67 pM cHET, and 10.57 pM HET). Despite AmMP having a lower median at the Hatched time point, the trend of increased concentrations holds as the AmMP IQR at Hatched (0.67-4.86 pM) was greater than the Incubation time point (0.58-2.07 pM).

Within the SW a similar temporal trend in dTRC concentrations was observed to the HZ where the dTRC concentrations at Spawn were frequently the highest measured (Figure 3). The median concentrations of all dTRC increased from Pre-Spawn (0.21 pM B1, 1.89 pM HMP, 7.76 pM AmMP, 70.57 pM cHET, and 1.47 pM HET) to Spawn (0.33 pM B1, 2.56 pM HMP, 12.50 pM AmMP, 103.27 pM cHET, and 10.84 pM HET). The median concentrations of all dTRCs except cHET decreased from Spawn to Incubation (0.23 pM B1, 2.26 pM HMP, 83.98 pM AmMP, 166.67 pM cHET, and 3.14 pM HET). All dTRCs saw a decrease in median concentration from the Incubation to Hatched time points (0.22 pM B1, 0.87 pM HMP, 6.45 pM AmMP, 44.13 pM cHET, and 1.02 pM HET).

### Relationship between dTRCs, microbial community diversity measures, and abiotic parameters

Spearman correlations between SW and HZ dTRC concentrations (all correlations significant unless indicated otherwise) were unique to each of these sample types. B1 concentrations in the HZ negatively correlated with HMP, cHET, and HET concentrations and showed positive correlations with AmMP (Figure 4A). In the HZ the biosynthetic precursors HMP and cHET positively correlated, yet the degradation products, HET and AmMP negatively correlated (Figure 4A). SW B1 concentrations correlated positively with HMP, HET, AmMP (Figure 4B). No significant negative co-correlations existed between dTRCs in the SW (Figure 4B). SW pH negatively correlated with concentrations of B1, HMP, and AmMP (Figure 4B).

**Figure 4.**
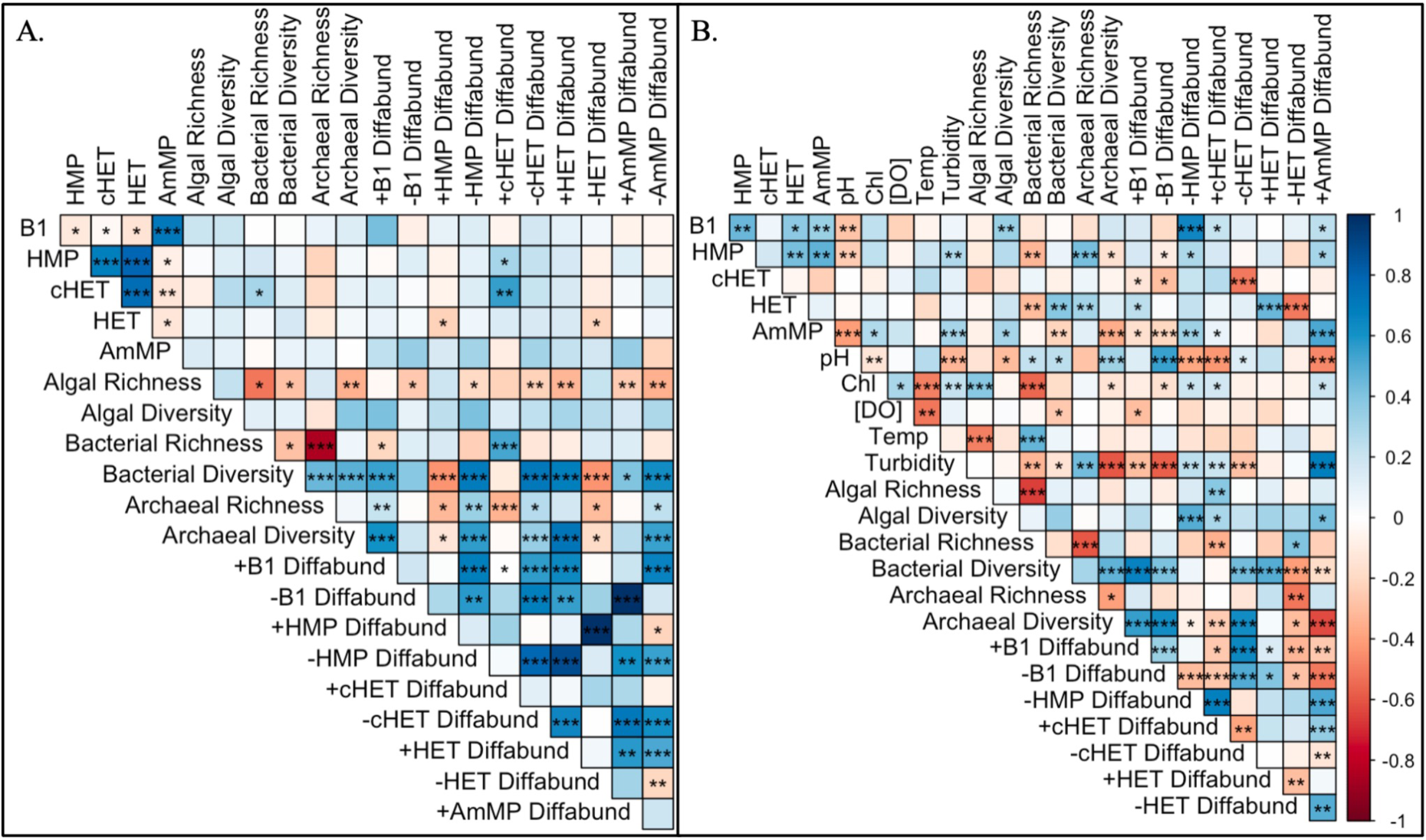
Correlogram of transformed and normalized A. HZ and B. SW dTRC concentrations, bacterial, archaeal, and algal (eukaryotic algae and Cyanobacteria) ASV richness and Shannon diversity, and differentially abundant ASV richness with each dTRC. “+” and “-” dTRC “Diffabund” indicates ASVs that are display positive and negative LFCs with dTRC concentrations, respectively. Positive correlations are depicted in shades of blue while negative correlations are depicted in shades of red. Stars indicate statistically significant correlative relationships (p < 0.05).

Other SW abiotic factors including temperature, chlorophyll-a, and dissolved oxygen concentrations showed no significant correlations with dTRC concentrations, and turbidity positively correlated with HMP and AmMP concentrations (Figure 4B). HZ bacterial richness positively correlated with cHET concentrations, and no other HZ microbial diversity matrices (richness and Shannon diversity) displayed significant associations with dTRC concentrations (Figure 4A). In SW samples algal Shannon diversity correlated positively with B1 and AmMP concentrations, yet algal richness and chlorophyll-a did not display this trend (Figure 4B).

Bacterial SW richness negatively correlated with HMP and HET concentrations and bacterial SW Shannon diversity positively correlated with HET and negatively correlated with AmMP. Archaeal SW richness positively correlated with HMP and HET concentrations and archaeal SW Shannon diversity negatively correlated with HMP and AmMP concentrations (Figure 4B). Notably, chlorophyll-a positively correlated with algal richness but not algal Shannon diversity.

Microbial richness, Shannon diversity, and the richness of differentially abundant ASVs correlated with dTRC concentrations to a higher degree in the WC than the HZ (Figure 4). The direction of the Log-Fold Change (LFC; + or - direction) from differential abundance results displayed in Figure 5 generally did not inform the direction of Spearman correlations (+ or -) between the richness of differentially abundant ASVs and dTRC concentrations, though some exceptions to this trend were present. In both sample types the richness of +B1 differentially abundant ASVs correlated positively with B1 concentrations, and the opposite was true for -B1 differentially abundant ASVs, yet none of these correlations were significant (Figure 4). The richness of SW -HMP, +cHET, and +AmMP differentially abundant ASVs all positively and significantly correlated with B1 concentrations (Figure 4B). Significant Spearman correlations where the LFC direction of differentially abundant ASVs (Figure 5) matched the Spearman correlation direction between differentially abundant ASV richness and dTRC concentrations included SW -cHET, +HET, -HET, and +AmMP (Figure 4B) and HZ +cHET and -HET (Figure 4A).

**Figure 5.**
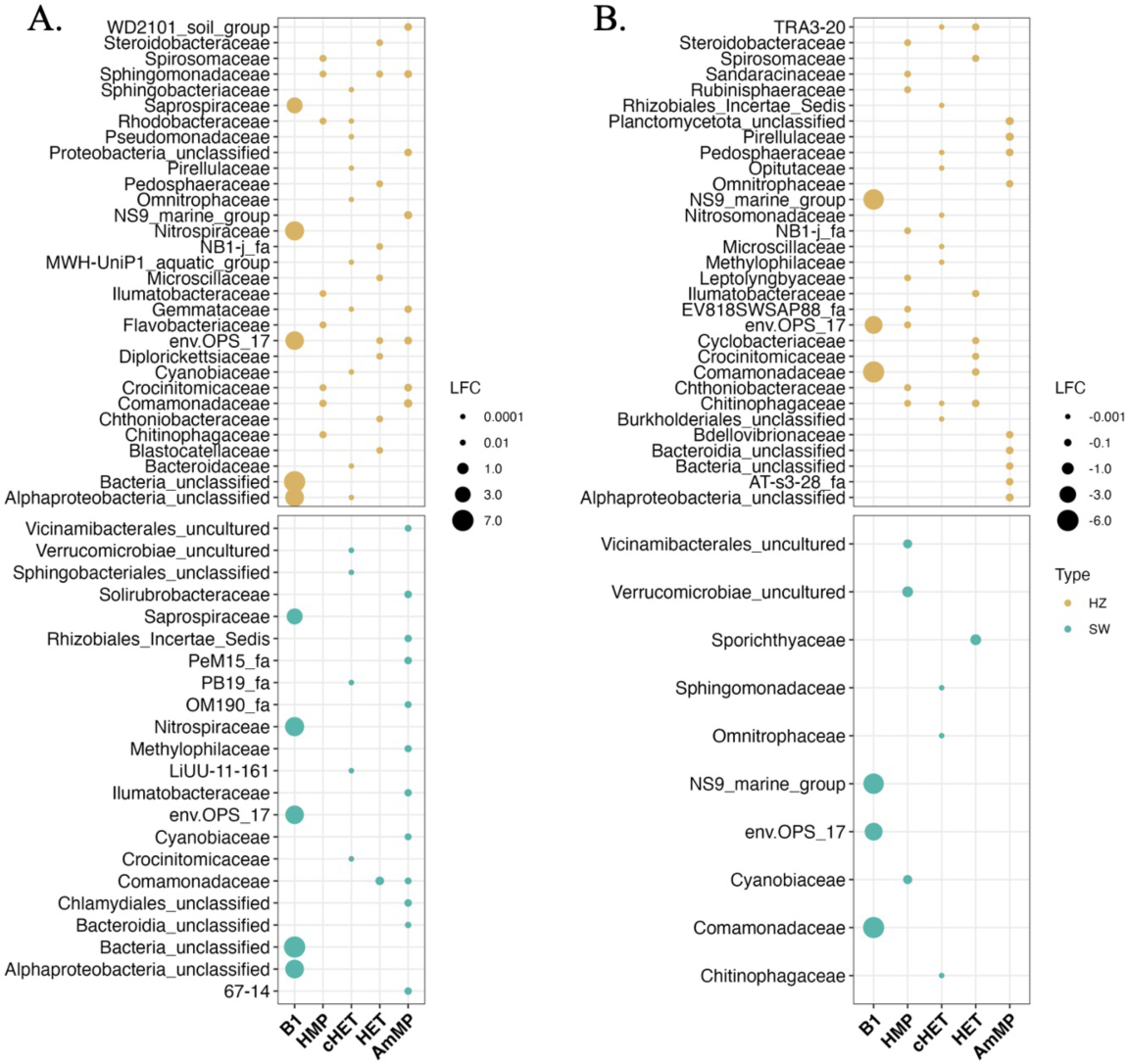
Bubble plot of A. positive and B. negative LFCs of differentially abundant ASVs. ASVs are significantly associated (p < 0.05) with raw dTRC concentration values. Each y-axis family represents the top 10 most differentially abundant ASVs per dTRC. Bubble color corresponds to sample type and the sizes of bubbles is related to LFC magnitudes.

Abiotic factors greatly influenced bacterial, archaeal, and algal richness and Shannon diversity in the SW. pH negatively correlated with algal Shannon diversity yet positively correlated with bacterial richness and bacterial and archaeal Shannon diversity (Figure 4B). Dissolved oxygen and temperature negatively correlated with bacterial Shannon diversity and bacterial richness, respectively (Figure 4B). Chlorophyll-a correlated positively with algal richness and negatively with bacterial richness and archaeal Shannon diversity (Figure 4B).

Turbidity correlated with bacterial and archaeal, yet not algal, richness and Shannon diversity. This parameter positively correlated with archaeal richness and negatively correlated with bacterial and archaeal Shannon diversity and bacterial richness.

### Sacramento River microbial communities differ by diversity and sample site characteristics

HZ bacterial, archaeal, and algal communities displayed higher diversity (all diversity measured with the Shannon diversity index) than those of the SW in all Sacramento River watershed regions (Figure S2). SW Battle and Butte Creek samples displayed the highest microbial diversity, followed by the Feather River, Clear Creek, and Sacramento River (Figure 6A). HZ diversity was similarly higher in Battle and Butte Creeks than all other samples and the Feather River showed the lowest HZ diversity. Clear Creek and Sacramento samples displayed nearly identical medians (Clear Creek = 6.26; Sacramento = 6.25) in HZ diversity. Temporal trends in diversity were less apparent but displayed a slight decrease in medians across time points (Pre-Spawn = 6.20; Spawn = 5.80; Incubation = 5.88; Hatched = 5.17; Figure 6B). Results from Kruskal Wallis tests indicated HZ sample diversity significantly differed by time point (p = 0.014), yet this was not true for SW samples (p > 0.05). SW samples significantly differed by region (p = 0.009) and HZ sample diversity did not significantly differ by region (p > 0.05).

**Figure 6.**
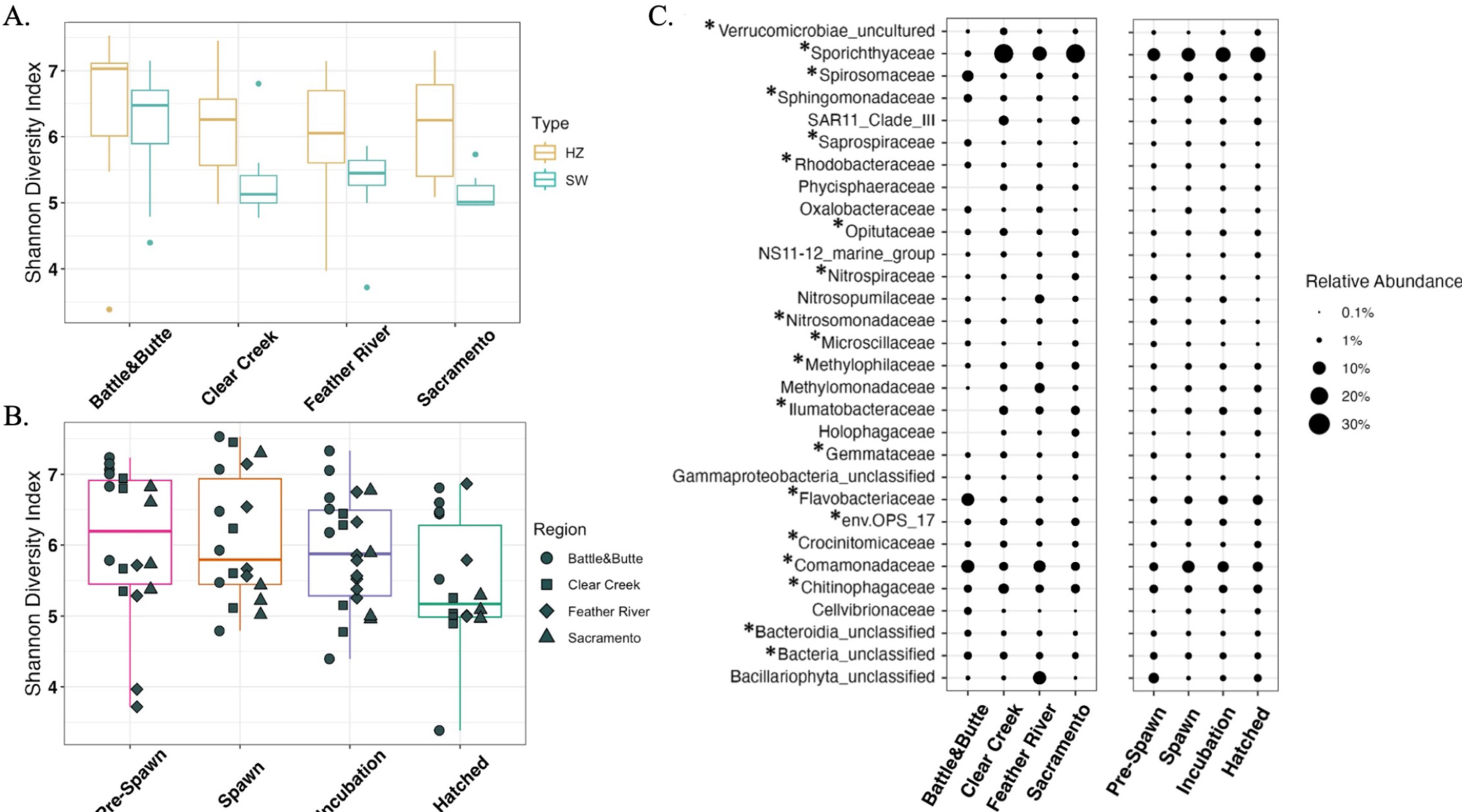
Microbial community diversity. Shannon diversity index values of bacterial, archaeal, and eukaryotic algal communities of SW and HZ samples binned by A. region and B. time point, and C. relative abundance of ASVs displaying the top 30 most abundant families by region and time point. Boxes in A. are colored by sample type and in B. by time point. Bubble size in C. represents relative abundance of specific ASVs and asterisks next to families represent those that contain species found to be differentially abundant with at least one dTRC.

Total rarefied bacterial and archaeal communities significantly differed by region, time point, and sample type (CAP PERMANOVA p = 0.001, 0.018, and 0.003, respectively). CAP1 explained 22.1% of variance and CAP2 explained 7.8% of variance (Figure S2). Significant differences in the variance of communities representing sample sites of each region also existed (permutest p = 0.0001), but not when samples were grouped by time point or sample type.

Within-region variance was also highest in Feather River samples, based on the centroid size of these sites on the CAP ordination (Figure S2). The bacterial and archaeal communities of Battle and Butte Creek differed greatly in composition from all other samples, especially on CAP1, and displayed low within-region variance (Figure S2).

### Bacterial, archaeal, and algal community compositions are unique by sample type

The total rarefied community of both sample types included 17,733 bacterial, archaeal, and eukaryotic algal ASVs. A small number of abundant phyla dominated in relative abundances across both sample types and represented taxa from all three domains. These included Proteobacteria (SW: 31.66%; HZ: 33.68%), Bacteroidota (SW: 22.46%; HZ: 19.86%), Actinobacteriota (SW: 20.16%; HZ: 13.75%), Planctomycetota (SW: 4.27%; HZ: 6.09%), Verrucomicrobiota (SW: 4.60%; HZ: 5.71%), Acidobacteriota (SW: 2.33%; HZ: 3.61%), Ochrophyta (SW: 2.64%; HZ: 3.16%), Unclassified Bacteria (SW: 1.77%; HZ: 2.43%), Crenarchaeota (SW: 1.82%; HZ: 1.44%), and Bdellovibrionota (SW: 1.40%; HZ: 1.54%). Notably, a majority of the top most abundant families across sample sites were differentially abundant with at least one dTRC (20/30; Figure 6C).

Feather River SW samples were significantly influenced (each vector p < 0.05) by water turbidity and chlorophyll-a concentrations. The concentration of dissolved oxygen significantly influenced the placement of SW Clear Creek and Sacramento sites and total dissolved solids and pH influenced both Battle and Butte Creek and Sacramento River samples (Figure 7A). dTRC concentrations did not significantly influence (each vector p > 0.05) the placement of any samples of either ordination (lack of dTRC vectors on Figure 7). Significant differences existed between SW communities by region (PERMANOVA p = 0.001) but not by time point (p > 0.05). Significant differences in within-group variance existed when SW samples were grouped by region (permutest p = 0.001) but not when SW samples were grouped by time point (p > 0.05).

**Figure 7.**
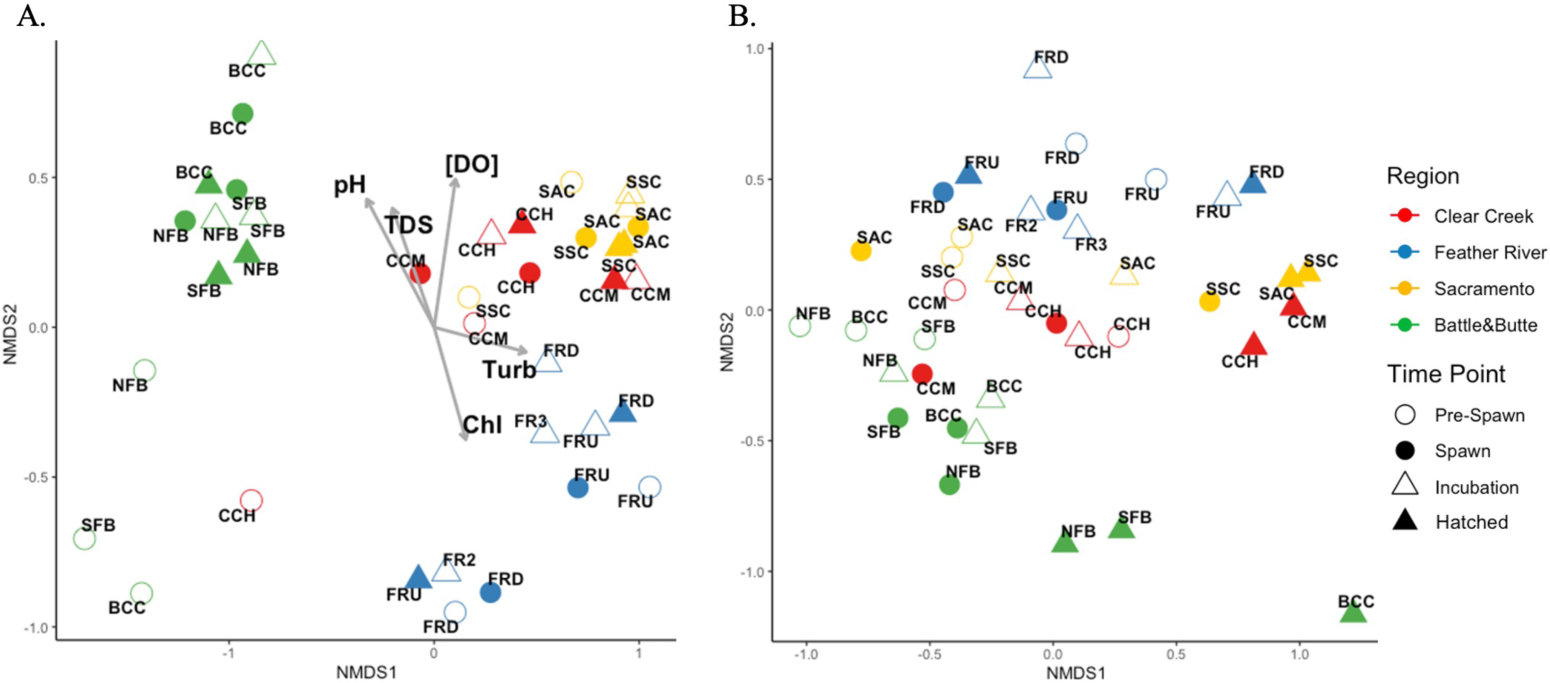
Bray-Curtis dissimilarity NMDS plots of A. SW (stress = 0.1082) and B. HZ (stress = 0.1219) bacterial, archaeal, and eukaryotic algal communities. Vectors display significant (p < 0.05) correlations between abiotic factors measured in the SW and sample site placements on the ordination. Points on the ordination are labeled by station, colored by region, and shapes correspond to time point. [DO] = Dissolved Oxygen (mg/L); Chl = Chlorophyll-a; TDS = Total Dissolved Solids; Turb = Turbidity.

Significant differences existed between HZ samples by both region (PERMANOVA p = 0.001) and time point (PERMANOVA p = 0.011), and no significant differences existed between within-group variance of either variable in the HZ (permutest p > 0.05). Battle and Butte Creek samples displayed similarities in community compositions regardless of both sample type and time point (Figure 7). However, no discernable patterns were observed for SW sample communities when grouped by time point (Figure 7A). In particular, the Sacramento River stations, SSC and SAC clustered closely together independent of time point (Figure 7A). All Battle and Butte Creek HZ microbial communities displayed high similarity by each of the time points and Clear Creek and Sacramento samples displayed similarities during the Hatched time point (Figure 7B). Mantel test results indicated both WC and HZ sample community dissimilarity significantly increased as a function of increased geographic (Haversine) distance between sample sites (SW: Mantel test statistic = 0.272; p = 0.0002; HZ: Mantel test statistic = 0.241; p = 0.0002). Community dissimilarity was not significantly associated with dTRC concentrations (p > 0.05) in the HZ. SW community dissimilarity was significantly associated with SW HMP concentrations (Mantel test statistic = 0.233; p = 0.017) and, to a lesser degree, with HET concentrations though this was not significant (Mantel test statistic = 0.173; p = 0.065).

### ASV relative and differential abundances are linked to dTRC concentrations

ANCOM-BC results indicated associations (all significant: p < 0.05) between ASVs in SW and HZ samples and every dTRC, especially B1 (+ and – LFCs; Figure 5). Families of differentially abundant ASVs in positive and negative LFC associations with B1 exhibited similar relative abundances between SW and hyporheic samples, and included Saprospiraceae (SW: 1.06%; HZ: 0.77%), Nitrospiraceae (SW: 0.78%; HZ: 0.91%), ENV OPS-17 (Sphingobacteriales Order; SW: 2.18%; HZ: 2.14%), Unclassified Bacteria (SW: 1.77%; HZ: 1.82%), Unclassified Alphaproteobacteria (SW: 0.41%; HZ: 0.52%), NS9 marine group (Flavobacteriales order; SW: 0.08%; HZ: 0.12%), and Comamonadaceae (SW: 6.88%; HZ: 5.71%) (Figure 6C).

Thiamine biosynthetic precursor compounds (cHET and HMP) and degradation products (HET and AmMP) showed much lower LFC magnitudes than B1 (Figure 5). Families of ASVs that were both differentially abundant in the HZ and with dTRC concentrations included Nitrospiraceae, Blastocatellaceae, Methylophilaceae, Uncultured Bacteria, Pedosphaeraceae, Rhizobiales *incertae sedis*, Chitinophagaceae, and Comamonadaceae (Figure S3; Figure 5). Only two families of ASVs, Saprospiraceae and Leptolyngbyaceae, were both differentially abundant in the SW and with dTRC concentrations (Figure S3; Figure 5). A sum of 342 ASVs were differentially abundant in association with each dTRC in HZ samples and a sum of 44 ASVs were differentially abundant in association with each dTRC in SW samples (grouped into each Figure 5 y-axis family).

The relative abundance of ANCOM-BC-identified ASVs displayed unique trends in samples grouped by region and binned dTRC concentrations (see methods). Samples grouped into the low B1 bin contained high relative abundances of Flavobacteraceae (4.45%) compared to Flavobacteraceae in normal (3.69%) and high B1 bins (3.07%), though all three bins still displayed high relative abundances of Flavobacteraceae. All other differentially abundant taxa displayed numerically even relative abundances across B1 concentration bins. Sporichthyaceae and Ilumatobacteraceae ASVs spiked in relative abundance in samples grouped into the low HET bin (17.56% and 3.19%, respectively) and were much lower in relative abundance in the normal (8.90% and 1.47%, respectively) and high (7.99% and 1.33%, respectively) HET bins. Similar to HET, the low HMP bin spiked in the relative abundance of Sporichthyaceae (16.19%) ASVs, whose relative abundance decreased in normal (12.38%) and high HMP (7.88%) bins. Opposingly, Sporichthyaceae ASVs spiked in relative abundance in samples grouped into the high AmMP bin (16.75%) and decreased in relative abundance in the normal (11.71%), and low (8.65%) AmMP bin.

Clear regional trends in the relative abundances of families of ANCOM-BC-identified ASVs were apparent. Butte and Battle Creek samples contained high relative abundances of Flavobacteriaceae, Spirosomaceae, and Comamonadaceae ASVs (8.58%, 6.47%, and 9.06%, respectively; Figure 6C) compared to the Feather River (1.86%, 1.49%, and 7.39%, respectively; Figure 6C), Clear Creek (1.43%, 1.26%, and 3.49%, respectively; Figure 6C), and Sacramento (1.11%, 1.35%, and 3.59%, respectively; Figure 6C). The opposite was true for Ilumatobacteraceae and Sporichthyaceae, which displayed high relative abundances in the Feather River (2.59% and 10.84%, respectively; Figure 6C), Clear Creek (3.37% and 21.29%, respectively; Figure 6C), and Sacramento River (3.44% and 20.93%, respectively; Figure 6C) compared to Battle and Butte Creek samples (0.05% and 1.36%, respectively; Figure 6C).

Though diatoms were not identified as being differentially abundant with dTRC concentrations, it is notable that Feather River sample sites contained high relative abundances of Unclassified Bacillariophyta ASVs (diatoms; 8.39%) and these ASVs were found in relative abundances below 1% in all other regions (Figure 6C). SAR11 clade III and CL500-3 (Phycisphaeraceae) were also higher in relative abundances in Sacramento River, Feather River, and Clear Creek samples than Battle and Butte Creek samples (Figure 6C) and in low HET and HMP bins than normal and high bins. SAR11 clade III taxa had relative abundances of 2.54%, 1.11%, and 1.00% in low, normal and high HET bins, respectively and CL500-3 taxa had relative abundances of 1.23%, 0.88%, and 0.69% in low, normal, and high HMP bins, respectively.

## DISCUSSION

The association between microbial communities and concentrations of dTRCs has not been investigated in freshwater systems despite an increase in reported incidence of TDC in these environments (1, 68). We determined the spatial and temporal distributions of dTRCs and the related microbial communities in the Sacramento River watershed in the SW and HZ (Figure 1). Environmental thiamine chemistry is a new and rapidly developing field where much remains to be discovered. Simultaneous, direct dTRC measurements have never been conducted in freshwater systems and yet such an approach is the best way to obtain a synoptic understanding of availability of this key vitamin. Thiamine has been infrequently measured previously in aquatic and estuarine systems (33–35). Of these studies, LCMS based thiamine measurements have only been conducted in the Moulouya River, Africa where mean thiamine concentrations were observed to be 4.5 pM (35). Distributions of dTRCs in marine systems have been studied more extensively, yet only roughly 500 discrete measurements have been published. A recent report from the North Atlantic observed dTRC concentrations to be within the picomolar range (7.35–353 pM for B1, 0.09–3.45 pM for HMP, 1.76–113.5 pM for AmMP, 0.03–30.8 pM for HET, and 3.76–145 pM for cHET) (26). Rate-based measurements of thiamine cycling (e.g., biological uptake, excretion, abiotic degradation) are even more sparse due to the complexity of the assays, which makes the interpretation of the available standing-stock dTRC concentrations challenging (69). Measured concentrations of dTRCs in environments are based on the equilibrium between microbial biosynthesis, dTRC excretion from cells due to lysis by viruses or grazers, microbial uptake of TRCs from the dissolved pool, and dTRC degradation by temperature, pH, UV light, and enzymatic activity (70). For this study we used an inference-based approach, pairing microbial community composition analysis with dTRC measurements to investigate the role of river and HZ microbial communities in the cycling of dTRCs, with the ultimate goal of making linkages to thiamine deficiency complex in Chinook salmon.

### Thiamine cycling differs in marine and freshwater systems

The differences between the observed dTRC concentrations in the Sacramento River and those previously reported in marine systems indicate that the thiamine cycle in each environment is controlled by a unique set of drivers. B1 concentrations observed in the Sacramento River SW and HZ (Figure 2, Table 1) are between two and four orders of magnitude lower than what has been reported in marine environments where the interquartile range of SW concentrations of B1 range between 15-64 pM and the interquartile range of sediment porewater B1 concentrations range between 237-462 pM (26, 43). Marine porewater B1 concentrations have only been measured in the deep ocean sediments (890 m water depth) and samples were collected over the complete redox zonation ranging from oxic to anoxic (43). The HZ samples in this study are from unconsolidated, oxic, river gravels, thus the comparison between these sample types is challenging. The interquartile concentration ranges of both the pyrimidine compounds HMP and AmMP from this study (Figure 2, Table 1) overlap those previously reported in the marine environment which are 0.3-5.1 pM HMP and 5.3-12.1 pM AmMP (26, 38, 42, 48). The thiazole moieties cHET and HET are both reported to be present at lower concentrations in the marine environment than in the Sacramento River SW where median concentration values were 8.4 and 11.3 times higher respectively (26)(Table 1). The authors are not aware of any published marine or freshwater sediment-associated HMP, AmMP, cHET, or HET concentrations.

It is unclear what factors drive the substantial differences between marine and freshwater SW dTRC concentrations. Physical and chemical differences between oceans and rivers could contribute to the observed concentration disparities as thiamine is known to abiotically degrade at high pH levels and with UV light (71–73), which could impact river SWs more readily than the marine water column. Additionally, the differences in mixing and advective processes between rivers and the ocean could have a major impact on dTRC concentrations. Further, the role of the distinct microbial communities in marine and freshwater environments cannot be discounted as biological activity (e.g., dTRC uptake and production) contributes to the standing stocks of each of these compounds in each environment. The disparities that we observed between marine and freshwater dTRC concentrations suggest that findings from investigations on thiamine cycling in marine ecosystems could be poor predictors of freshwater thiamine dynamics. Future research will be required to fully understand the factors controlling thiamine cycling in marine and freshwater systems.

### dTRC distributions in the Sacramento River system

All dTRCs were detected in both the SW and HZ of the Sacramento River system (Figure 2, Tables 1, S1). This is the first report of the distribution of all these compounds in aquatic environments. The concentrations of B1 across the entire watershed were exceedingly low, with median concentrations in the sub-picomolar range (0.21 pM HZ, 0.23 pM SW). B1 concentrations were the lowest of the dTRCs measured in this study (Figure 2, Table 1). We hypothesize that B1’s low concentration is due to a combination of its abiotic instability and its biological importance, which would drive rapid B1 uptake (microbial and metazoan) from the extracellular environment. Microbial community results support this hypothesis because differentially abundant ASVs had more measurable impacts on B1 than all other dTRCs regardless of sample type (LFC magnitudes in Figure 5). The known cellular enzymatic half saturation constants for thiamine are in the nanomolar range and indicate that cells must have high affinity uptake mechanisms for this coenzyme (74). Thiamine auxotrophy (i.e., obligate requirement) is known to be prevalent in many groups of abundant and environmentally important microorganisms which creates a robust demand for dissolved thiamine and its congeners in most environments (22, 37, 42). Additionally, we observed that the B1 degradation product AmMP was present at concentrations between one and two orders of magnitude higher than B1. These data taken together suggest that microbial activity coupled with abiotic degradation could be responsible for driving equilibrium B1 concentrations (e.g., balance between production, uptake, and degradation) into the femtomolar range.

The measured concentrations of HMP and cHET which are the biosynthetic precursors for B1, are between 2 and three orders of magnitude greater than B1 (Figure 2, Table 1). These data do not support the alternate hypothesis that low B1 concentrations are due to low rates of production, and instead provide substantial evidence for relatively high rates of thiamine biosynthesis in both the SW and the HZ. The sole known source for HMP and cHET is the thiamine biosynthesis pathway (25), therefore it is somewhat surprising to see these intracellular tracers of the thiamine biosynthesis pathway in the extracellular dissolved pool. However, multiple studies have reported these compounds to be present in spent culture media and the environment, and have implicated cellular leakage, excretion, cellular lysis, and sloppy feeding as mechanisms for their release (26, 37, 42, 70). Our data paint a picture of a dynamic thiamine cycle in the Sacramento River system where relatively high rates of microbial thiamine biosynthesis, as a function of microbial community structure, are balanced with the removal processes of abiotic degradation and microbial (or metazoan) uptake resulting in sub-picomolar dissolved thiamine concentrations. Future investigations will be required to quantify the rates that each dTRC are produced and removed to fully constrain the freshwater thiamine cycle.

We observed that biosynthetic precursors of thiamine, HMP and cHET, are in higher concentrations in the HZ than in the overlying SW which we interpret to be evidence for active thiamine biosynthesis in the HZ (Figure 2). This observation provides the strongest evidence to date that the HZ is source of dTRCs to the SW, building on previous from marine and freshwater systems (35, 43). Riverine HZs are a major source of microbes and metabolites to the overlying river, and the residence time of the water in the river is directly proportional to the abundance and diversity of both microbes and metabolites (75, 76). It is not clear how variations in river flow influence dTRC flux from the sediments, but changes in HZ residence time likely influences this process. Additional abiotic and biotic factors such as organic carbon loading, sediment particle size, and discharge rate, which impacts permeability and the extent of the HZ and its exchange with the SW, respectively (77), likely influence dTRC SW concentrations in ways beyond the scope of this study. Our data point to a robust benthic-associated source of dTRCs to the overlying SW. These findings are of special interest, as they suggest a substantial naturally occurring source of thiamine in spawning gravels which could serve to supplement early life stages of salmon that are thiamine deficient.

### Regional variability in dTRC concentrations

Despite observed regional variations in river physical and chemical parameters, only weak regional differentiation in dTRC concentrations was observed (Figure 1, 2, S1). Sampling locations for this study were chosen to assess the impact of a gradient of anthropogenic impacts and salmon spawning success on dTRC availability. We initially hypothesized that due to these varying impacts, dTRC concentrations would vary based on river. This hypothesis was supported by our physical and chemical data from these rivers which demonstrate differing hydrological regimes in tributaries below high head dams and those with lower levels of anthropogenic impacts (Figure 1). For example, multiple precipitation events occurred during our sampling period which caused pulses of high flow in Battle, Butte, and Clear Creeks (Figure 1), however, these flow events were prevented by upstream dams at our sites in the Sacramento and Feather Rivers. The existence of the Oroville dam upstream of the Feather River sampling sites may have contributed to the strong regional signature observed at these sites, where the pH and temperature were consistently lower than other tributaries while the turbidity and chlorophyll-a were consistently higher (Figure 1). Spearman correlations showed that pH negatively correlated with SW B1, HMP, and AmMP concentrations, yet did not impact cHET and HET (Figure 4). The consistently lower pH at the Feather River sites, resulting in increased chemical stability of dTRCs, likely caused the slight observed increase in B1, HMP, and AmMP concentrations in the Feather River SW relative to other regions (Figure 1, S1). Kruskal-Wallis tests showed that B1 within the surface water differed significantly by region (p-value<0.05).

This regional increase in dTRC concentrations may have been limited by the higher abundances of potentially-auxotrophic algae at this site as evidenced by higher chlorophyll-a concentrations and diatom ASV relative abundances (Figure 1, Figure 6C)(40). These observations demonstrate the complexity of thiamine cycling, where multiple variables impact both production and removal processes of dTRCs.

### Temporal influences on dTRC concentrations

Strong temporal patterns of dTRC concentrations across the sample sites were observed which suggest that there could be linkages between microbial dTRC production and the Chinook salmon life cycle (Figure 3). There was a clear seasonal signature observed in our chemical data where temperature across all sample sites decreased from September to January (Figure 1).

Additionally, we observed that chlorophyll-a and dissolved oxygen increased over this same time period (Figure 1). These trends reflect the seasonal progression from late summer to mid-winter. However, when the dTRC concentrations are binned by sample time point they show a temporal pattern for all dTRCs that is decoupled from seasonality (Figure 3). We observed that maximum dTRC concentrations generally occurred at the Spawn (November) time point which occurred while the fall run Chinook salmon were actively spawning in the Sacramento River watershed (Figure 3). The Spawn maximum was generally followed by a decrease in dTRC concentration at the Incubation (December) time point which occurred while the fall run Chinook salmon eggs were incubating in the river gravels (Figure 3). At the final sample point Hatched (January), we assume that most fall-run Chinook salmon eggs that were in the gravel at the Incubation time point would have hatched and it is likely that the juveniles were still benthically associated (47). At Hatched in the HZ dTRC concentrations increased, while in the SW they decreased (Figure 3). Kruskal-Wallis tests showed that these temporal differences were statistically significant for HMP in the SW and AmMP and cHET in both the SW and HZ (p-value<0.05). The processes driving the observed temporal trend in dTRC concentrations across our sample sites are not clear, however our data do indicate that these processes are decoupled from the seasonal changes occurring in water chemistry (Figure 1C). Research has indicated that spikes in dTRC concentrations in marine environments are frequently related to shifts in the microbial community composition and activity (26, 38). In this study we observed that microbial communities in the HZ differ by sample time point, as opposed to the SW (Figure 7). Thiamine is a biologically valuable compound, and as such dTRC concentrations are frequently at a delicate equilibrium governed by the balance between production and removal processes. In order for a temporal trend to be observed over multiple heterogeneous tributaries across a large river basin, there must be a universal factor causing the shift. As the trends we observe in dTRC concentration are not linked to seasonal changes, we must look elsewhere for this missing variable.

### Human impacts on rivers influence microbial community compositions

The compositions of Sacramento River watershed microbial communities showed small-scale homogeneity and large-scale heterogeneity. The diversity of microbial communities can be dependent on both river reach (78, 79) and sample type (80). High similarity between microbial communities of stations in the same river region were observed and indicated a lack of evidence for river reach driving community heterogeneity (Figure S2). Regions with high anthropogenic impact (e.g., Sacramento, Feather River, and Clear Creek) also displayed more within-region variance than Butte and Battle Creek sites (Figure S2), providing evidence that anthropogenic impacts could increase microbial community dissimilarity between locations along the same river, and between the surface water and hyporheic zone. This could result in sporadic compositions of microbial communities along the spectrum of human-impacted rivers where Chinook salmon spawn. Human-constructed dams and the resulting reservoirs impede river flow and can alter microbial diversity in fluvial systems (80); Jones 2020; Maavara 2020}. Though our results do not contain above- and below-reservoir sample locations, we did observe differences between sites downstream of dam-formed reservoirs and sites with lower human-impact. Battle and Butte Creek sites displayed higher microbial diversity in both sample types than heavily-impacted river sites (Figure 6A), which provides evidence that anthropogenic impacts limit microbial diversity in the Sacramento River watershed.

### Microbial diversity and community compositions influence dTRC concentrations uniquely by sample type

While SW Shannon diversity differed by river region, HZ Shannon diversity differed by time point and not region in a similar manner to dTRC concentrations. However, few significant correlations existed between HZ Shannon diversity and dTRC concentrations (Figure 4A). Together these results provide evidence that HZ microbial diversity changes in accordance with temporal variation but does not greatly influence dTRC concentrations. River HZs are environments of high chemical complexity and microbially-driven biochemical and respiratory activity (81). High-level microbial diversity therefore may not reflect the biochemical complexity of this environmental system whereas the relative abundance of certain groups of ASVs could influence small-scale biochemistry, and therefore dTRC concentrations.

Contrary to the HZ, microbial diversity appears to greatly influence the availability of dTRCs in the SW (Figure 4B). Far more significant correlations were observed between SW microbial richness and Shannon diversity and dTRC concentrations, providing evidence that microbial diversity could be an indicator for dTRC availability in the SW. Microbial community compositions displayed regional signatures in both sample types (Haversine distance Mantel test results and Figures S2 & 7), yet only HZ samples display community compositions unique to salmon spawn time points. This finding provides evidence that the structure of microbial communities in the HZ, influenced by relative abundances of certain ASVs, could impact dTRC concentrations in occurrence with the Chinook salmon life cycle.

### Relationship between microbial taxa and dTRCs

Microbially-driven thiamine cycling is likely more complex in the HZ than water column based on the high numbers of differentially abundant ASVs with HZ dTRC concentrations, compared to those in the SW. Further, high concentrations of the thiamine biosynthetic precursor compounds, cHET and HMP in the HZ could signify thiamine biosynthetic activity from microbial synthesis and exudation. The larger LFC magnitudes of differentially abundant ASVs associated with B1, compared to the other dTRCs, gives evidence for intact thiamine availability being more highly influenced by the presence of certain taxa than the other dTRCs. The observation that a majority of highly relatively abundant taxa influence dTRC concentrations (Figure 6C) also suggests that abundant microbes impact SW and HZ dTRC availability to a high degree.

SW algal Shannon diversity influenced dissolved B1 availability to a higher degree than the relative abundance of particular algal taxa, algal richness, and Chlorophyll-a concentrations. Evidence for this finding was indicated by diatom ASVs being found in high relative abundances in Feather River samples, yet these taxa were not differentially abundant with any dTRC concentrations, though algal Shannon diversity significantly correlated with SW B1 concentrations (Figure 4B). These results follow previous findings, which have implicated the existence of algal blooms (low algal diversity states) with depleted levels of dissolved thiamine (40, 82). Conversely, individual bacterial taxa appeared to greatly influence dTRC concentrations. ASVs of several different families were found to be differentially abundant in both sample types and in both LFC directions with multiple dTRCs, especially B1, and included ENV OPS-17, Comamonadaceae, Unclassified bacteria (unclassified to phylum level), and NS9 marine group (Figure 5). ASVs within each of these families may therefore display site-specific thiamine-cycling characteristics influenced by unknown extracellular conditions.

Our data demonstrates human-impacted freshwater habitats harbor heightened relative abundances of bacteria that negatively influence thiamine availability than habitats with less human impact, which could influence the availability of B1 to Chinook salmon. Sporichthyaceae ASVs captured the above trend and the hgcl clade (genus annotation of Sporichthyaceae) was the most abundant genus across all samples sites yet were exceedingly low in relative abundances in Battle and Butte Creeks (Figure 6C). Hgcl clade taxa have previously been associated with eutrophic aquatic settings, which could correspond to diatom overabundances that could indicate algal bloom states (83). These ASVs also displayed heightened relative abundances in sample sites binned into low HMP and HET, yet no change in relative abundance in B1 bins, indicating evidence for precursor auxotrophy and drawdown. In addition to hgcI clade, several other genera exhibited higher relative abundances in high human-impact sites compared to Battle and Butte Creeks and included CL500-3 (Phycisphaeraceaein; Figure 6C), CL500-29 marine group (Ilumatobacteraceae; Figure 6C), and SAR11 clade III (Figure 6C). Each of these taxa displayed heightened relative abundances in low sample bins of HET and HMP. Further, CL500-3, CL500-29, and hgcI genera have been found in high relative abundances in reservoirs of past studies (84, 85). This provides limited evidence for human-made reservoirs seeding fluvial systems with bacteria that could draw down concentrations of certain dTRCs (HET and HMP).

Gene activity needs to be assessed to link the transcription (and lack of transcription) of thiamine biosynthesis and salvage genes to dTRC environmental concentrations. Though we were able to correlate ASV compositions with dTRC concentrations, the activity of thiamine-related gene pathways ultimately dictates the availability of these compounds to the environment through microbial production and consumption of thiamine-related compounds. Previous research in marine settings also show that microbial communities existing in structural disequilibria display heightened auxotrophy, limiting overall thiamine availability (26).

Anthropogenic impact is also known to limit the availability of thiamine to higher trophic organisms due to nutrient input and climate change factors (23). Linking human impact, which could increase microbial community disequilibria, with the transcription of thiamine-related genes and the concentrations of dTRCs in fluvial systems is therefore of future relevance to assess the drivers of thiamine availability in Chinook salmon spawning habitats with differing degrees of anthropogenic disturbances.

We observed that heavily human-impacted freshwater systems harbor microbial communities with diminished microbial diversity and unique community compositions, increased community dissimilarity between sites within the same river region, and higher relative abundances of taxa negatively associated with dTRC concentrations. River systems heavily impacted by human pollution have been previously shown to contain unique microbial community structures and diversity compared to pristine fluvial habitats (86). Several studies have also displayed the many negative health outcomes anthropogenic pressures, including pollutants, habitat loss, climate change, freshwater acidification, and the building of dams, have on salmon in marine and freshwater systems (87–90). Linking microbial community compositions and diversity to dTRC concentrations in rivers where salmon spawn that are heavily human-impacted is of great importance to obtain a comprehensive trophic view of TDC in natural fluvial systems. Though requiring more detailed research, the availability of thiamine to spawning Chinook salmon, incubating eggs, and hatched fry may therefore be constrained in fluvial settings impacted by human-made dams and other anthropogenic stressors.

## CONCLUSION

We hypothesized that there is a relationship between the presence of spawning fall run Chinook salmon and the availability of dTRCs in both the HZ and the overlying SW. We observed that the peak in dTRC abundance aligns with the presence of spawning salmon, which is followed by a decrease in concentration while early life stages of salmon are incubating in the river gravels. It is known that early life stages of salmon (eggs and embryos) can assimilate dissolved thiamine from their environment (21), therefore the decrease in observed dTRC concentration between the Spawn and Incubation time points could be caused by dTRC uptake by early life stages of salmon. Our understanding of aquatic thiamine dynamics are at an early stage of discovery, and additional research is required to complete our understanding of this system. However, the possible linkage between river HZ microbiomes and dTRC concentrations could point to a coevolutionary relationship where salmon spawn at times that take advantage of a seasonal increase in microbial thiamine production. Such a relationship between high dTRC concentrations and salmon spawning could serve to rescue eggs and embryos from maternally acquired thiamine deficiency complex. Alternative, and less evolutionarily complex, explanations for our observations also exist. For example, spawning salmon cause a substantial disturbance in the river gravels while digging redds (91). It is possible that this physical remodeling of the benthic environment causes a release of thiamine, promoting higher concentrations in the HZ and overlying SW. The decrease in dTRCs after spawning could be due to an increase in microbial activity and thus thiamine demand driven by increased organic carbon input from decaying salmon carcasses. It remains to be demonstrated that there is a causal link between salmon presence and dTRC availability, or that this pattern is repeatable over interannual cycles. Regardless, these data are an intriguing first step into understanding the thiamine dynamics of aquatic systems and their relationship with Chinook salmon.

## ACKNOWLEDGMENTS

This work was funded by the California Department of Fish and Wildlife grant #Q2196012. Additional personnel funding for CPS was provided by NSF grant DEB-1639033. Mass spectrometry instrumentation was supported by NIH grant 1S10RR022589-01. We thank Stephen Giovannoni for his support, mentorship, and usage of his laboratory space. We thank Jessica Büser-Young and Dale Honeyfield for their collegial reviews of this manuscript. We thank Jeff Morré, Elizabeth Brennan, Sarah Wolf, Chih-Ping Lee, and Matt Emard for their technical scientific support. Finally, we thank the broader community of Salmonid Thiamine Deficiency Complex scientists for helpful and inspirational conversations which have shaped the direction of this research.

## Data Availability Statement

The molecular datasets presented in this study can be found in the NCBI sequence read archive (92). PRJNA996220: https://www.ncbi.nlm.nih.gov/bioproject/996220. All dTRC data are published in Table S1.

