## Supplemental Information for "Connecting Thiamine Availability to the Microbial Community Composition in Chinook Salmon Spawning Habitats of the Sacramento River Basin"

**Running Title:**
Sacramento River Thiamine

**Suffridge, Christopher P.**[*+], Oregon State University, Department of Microbiology. 226 Nash Hall, Corvallis, OR 97331
**Shannon, Kelly C.**[*], Oregon State University, Department of Microbiology. 226 Nash Hall, Corvallis, OR 97331
**Matthews H.,** Oregon State University, Department of Microbiology. 226 Nash Hall, Corvallis, OR 97331
**Johnson R.,** NOAA Fisheries, Southwest Fisheries Science Center, Fisheries Ecology Division, 110 McAllister Way, Santa Cruz, CA 95060 and University of California Davis, Center for Watershed Sciences, 1 Shields Ave. Davis, CA 95616
**Jeffres C.,** University of California, Davis, Center for Watershed Sciences, One Shields Ave, Davis 95616
**Mantua N.,** NOAA Fisheries, Southwest Fisheries Science Center, Fisheries Ecology Division, 110 McAllister Way, Santa Cruz, CA 95060
**Ward, Abigail E.,** University of California, Davis, Center for Watershed Sciences, One Shields Ave, Davis 95616
**Holmes E.,** University of California, Davis, Center for Watershed Sciences, One Shields Ave, Davis 95616 AND California Department of Water Resources, 3500 Industrial Blvd. West Sacramento, CA 95691
**Kindopp, J.,** California Department of Water Resources, Division of Integrated Science and Engineering. 460 Glen Dr., Oroville, CA 95966.
**Aidoo M.,** Bronx Community College/CUNY, Bronx, NY 10453
**Colwell F.,** Oregon State University, College of Earth Ocean and Atmospheric Sciences; Oregon State University, Department of Microbiology. 226 Nash Hall, Corvallis, OR 97331

[*]Co-first authors

### SUPPLEMENTARY FIGURES AND TABLES

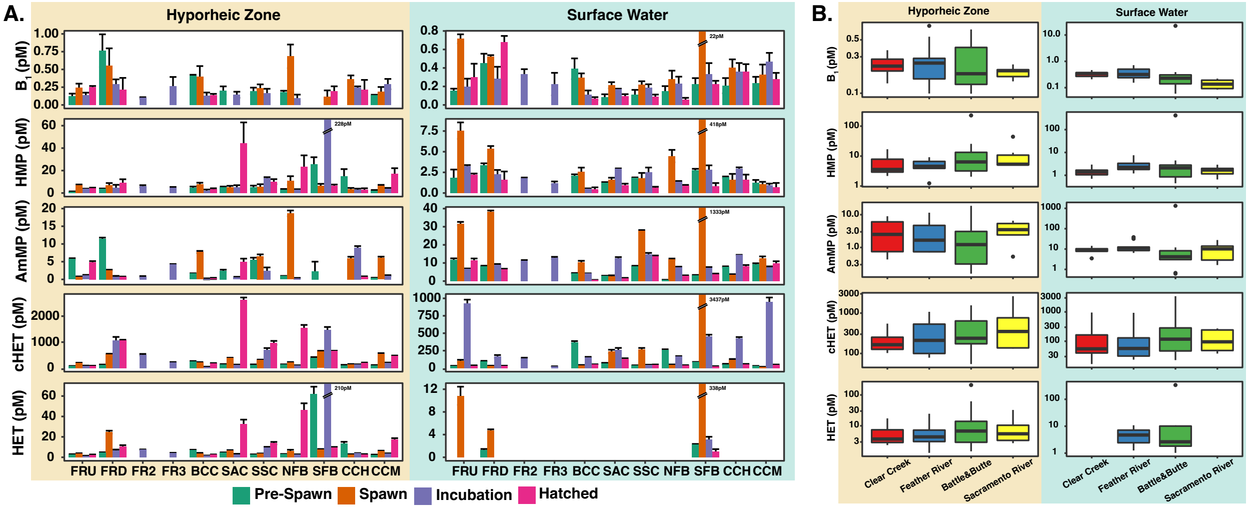


***Supp. Figure 1: Regional variations in dTRC concentrations.*** *A. Concentrations of dTRCs in the river surface water (blue) and hyporheic zone (tan) at each sampling location and time point. Station codes are described in Methods. Bar height and error bars represent the mean and standard deviation of three technical replicates. No bars indicate that dTRCs were not detected in the samples, except for FR2 and FR3 where samples were only collected at the In-Gravel time point. Full dataset is in Table S1. B. Boxplots of hyporheic zone (tan) and surface water (blue) dTRC concentrations are binned by region. Colors indicate tributary: Feather River (Blue), Sacramento River (Yellow), Clear Creek (Red), Battle and Butte Creeks (Green).*

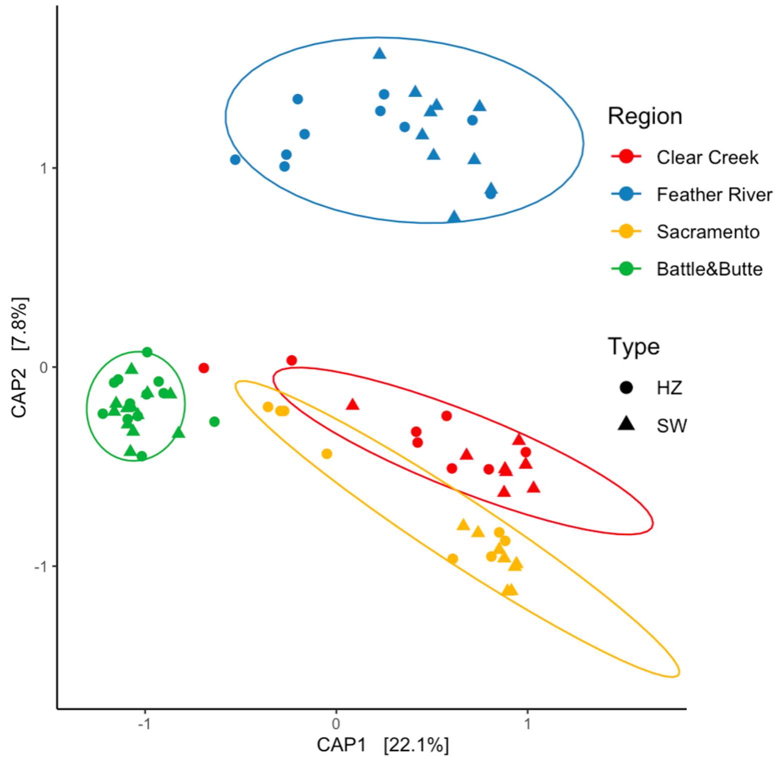


***Supp. Figure 2: Constrained-analysis of principal coordinates displaying differences in bacterial, archaeal, and algal communities.*** *Data are constrained by the variables: region, time point, and sample type. CAP1 described 21.5% of variance and CAP2 described 8.3% of variance. Colors correspond to region and shapes correspond to sample type.*

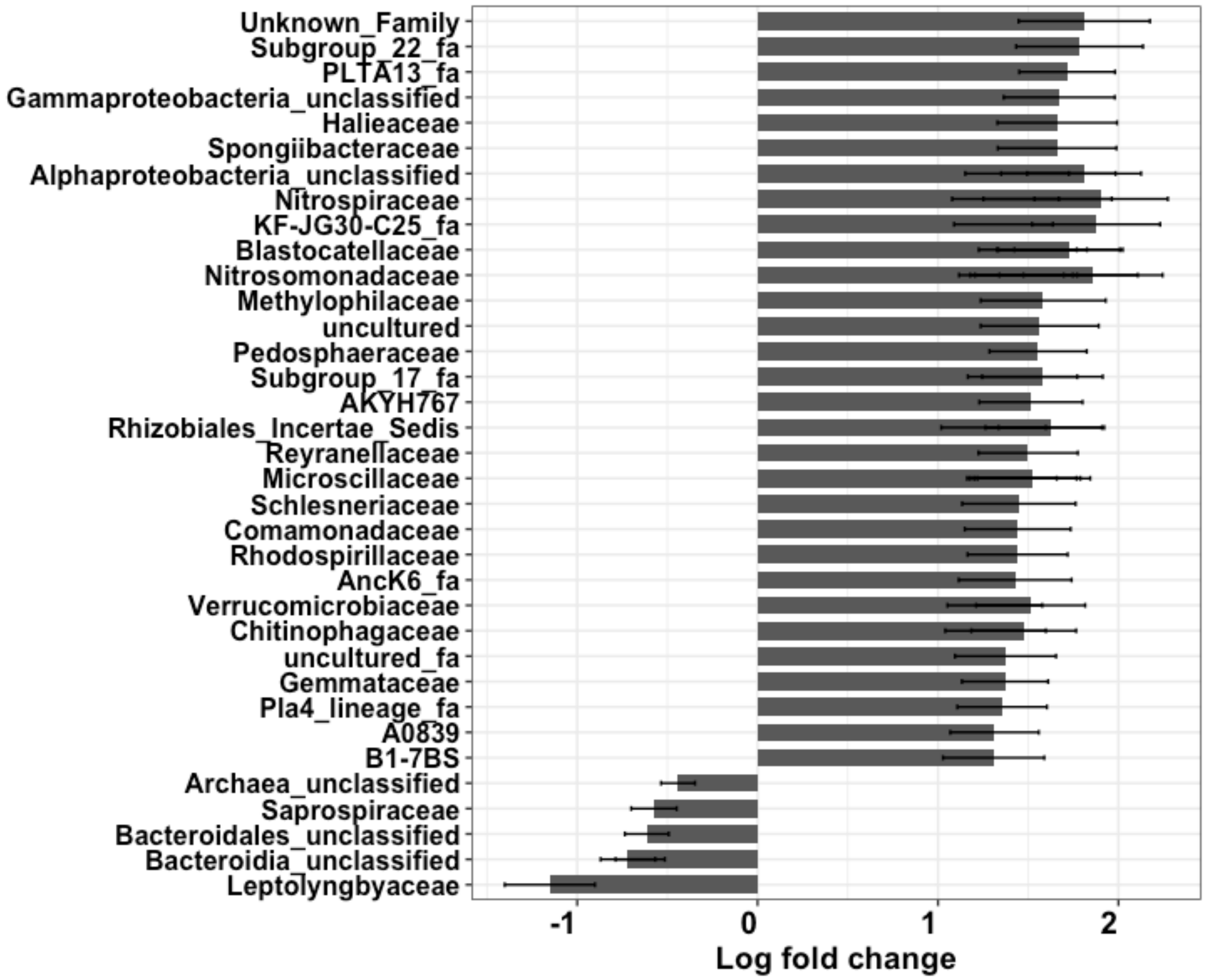


***Supp. Figure 3: Families of ASVs that are differentially abundant in hyporheic zone samples (+log-fold change) compared to surface water samples (-log-fold change).***

***Supp. Table 1: dTRC concentrations.*** *Values are means and standard deviations of three technical replicates. All concentrations are in picomolar. Blanks indicate compound was undetectable in the sample.*

| Time Point | Station | River | Type | B1 | HMP | AmMP | cHET | HET |
|---|---|---|---|---|---|---|---|---|
| Pre-Spawn | FRU | Feather River | HZ | 0.12 ± 0.04 | 1.24 ± 0.20 | 5.77 ± 0.21 | 78.67 ± 1.86 | 2.90 ± 0.15 |
| Pre-Spawn | FRD | Feather River | HZ | 0.76 ± 0.23 | 3.84 ± 0.20 | 11.50 ± 0.30 | 133.00 ± 4.36 | 4.63 ± 0.29 |
| Pre-Spawn | BCC | Battle&Butte | HZ | 0.41 ± 0.01 | 4.63 ± 1.14 | 1.63 ± 0.09 | 254.33 ± 16.92 | 7.28 ± 0.13 |
| Pre-Spawn | SAC | Sacramento River | HZ | 0.20 ± 0.06 | 5.12 ± 0.65 | 2.45 ± 0.29 | 139.33 ± 15.70 | 4.64 ± 0.14 |
| Pre-Spawn | SSC | Sacramento River | HZ | 0.20 ± 0.05 | 4.99 ± 1.62 | 5.51 ± 0.63 | 132.00 ± 10.58 | 2.74 ± 0.08 |
| Pre-Spawn | NFB | Battle&Butte | HZ | 0.18 ± 0.02 | 2.84 ± 0.76 | 0.91 ± 0.07 | 139.67 ± 3.51 | 2.97 ± 0.26 |
| Pre-Spawn | SFB | Battle&Butte | HZ | | 25.67 ± 6.23 | 2.26 ± 2.75 | 387.00 ± 36.01 | 62.20 ± 6.99 |
| Pre-Spawn | CCH | Clear Creek | HZ | | 14.93 ± 6.38 | | 102.60 ± 23.40 | 13.60 ± 1.57 |
| Pre-Spawn | CCM | Clear Creek | HZ | 0.13 ± 0.01 | 2.17 ± 0.50 | 0.43 ± 0.03 | 103.33 ± 14.57 | 2.37 ± 0.27 |
| Spawn | FRU | Feather River | HZ | 0.24 ± 0.06 | 6.87 ± 0.73 | 0.76 ± 0.08 | 201.33 ± 5.86 | 3.75 ± 0.20 |
| Spawn | FRD | Feather River | HZ | 0.55 ± 0.25 | 6.99 ± 1.88 | 2.57 ± 0.17 | 538.33 ± 16.62 | 24.60 ± 1.45 |
| Spawn | BCC | Battle&Butte | HZ | 0.40 ± 0.15 | 7.83 ± 1.36 | 7.73 ± 0.31 | 223.33 ± 7.51 | 3.93 ± 0.37 |
| Spawn | SAC | Sacramento River | HZ | | 5.27 ± 1.14 | | 391.67 ± 8.39 | 6.54 ± 0.51 |
| Spawn | SSC | Sacramento River | HZ | 0.24 ± 0.05 | 5.46 ± 2.37 | 6.55 ± 0.51 | 314.00 ± 13.86 | 3.57 ± 0.18 |
| Spawn | NFB | Battle&Butte | HZ | 0.68 ± 0.17 | 11.04 ± 3.95 | 18.63 ± 0.81 | 223.33 ± 10.26 | 6.29 ± 1.07 |
| Spawn | SFB | Battle&Butte | HZ | 0.12 ± 0.09 | 6.43 ± 1.76 | | 633.00 ± 33.05 | 7.65 ± 0.32 |
| Spawn | CCH | Clear Creek | HZ | 0.37 ± 0.05 | 3.29 ± 1.24 | 5.99 ± 0.41 | 139.67 ± 17.39 | 2.88 ± 0.19 |
| Spawn | CCM | Clear Creek | HZ | 0.19 ± 0.06 | 6.35 ± 0.91 | 6.06 ± 0.36 | 549.00 ± 27.73 | 6.06 ± 0.42 |
| Incubation | FRU | Feather River | HZ | 0.14 ± 0.04 | 3.76 ± 0.07 | 1.09 ± 0.19 | 90.90 ± 6.24 | 1.33 ± 0.22 |
| Incubation | FRD | Feather River | HZ | 0.30 ± 0.06 | 4.60 ± 2.72 | 0.73 ± 0.26 | 1061.33 ± 139.52 | 6.62 ± 0.95 |
| Incubation | FR2 | Feather River | HZ | 0.10 ± 0.01 | 5.98 ± 1.00 | 0.73 ± 0.19 | 516.33 ± 29.67 | 7.47 ± 0.22 |
| Incubation | FR3 | Feather River | HZ | 0.26 ± 0.13 | 4.54 ± 0.85 | 4.26 ± 0.08 | 221.67 ± 11.59 | 4.14 ± 0.27 |
| Incubation | BCC | Battle&Butte | HZ | 0.13 ± 0.04 | 2.05 ± 1.06 | 0.16 ± 0.13 | 54.73 ± 0.74 | 1.49 ± 0.23 |
| Incubation | SAC | Sacramento River | HZ | 0.14 ± 0.04 | 4.96 ± 2.01 | 0.53 ± 0.21 | 132.00 ± 4.36 | 2.84 ± 0.56 |
| Incubation | SSC | Sacramento River | HZ | 0.17 ± 0.06 | 12.97 ± 1.16 | 2.40 ± 0.97 | 705.33 ± 66.76 | 9.47 ± 0.74 |
| Incubation | NFB | Battle&Butte | HZ | 0.10 ± 0.05 | 2.88 ± 0.52 | 0.33 ± 0.09 | 99.73 ± 9.34 | 2.81 ± 0.12 |
| Incubation | SFB | Battle&Butte | HZ | | 227.67 ± 116.84 | | 1423.33 ± 185.02 | 209.67 ± 25.01 |
| Incubation | CCH | Clear Creek | HZ | 0.24 ± 0.02 | 2.95 ± 0.94 | 8.96 ± 0.44 | 135.67 ± 10.97 | 3.93 ± 0.18 |
| Incubation | CCM | Clear Creek | HZ | 0.29 ± 0.07 | 3.89 ± 0.65 | 1.05 ± 0.14 | 206.00 ± 8.00 | 3.55 ± 0.34 |
| Hatched | FRU | Feather River | HZ | 0.25 ± 0.01 | 4.04 ± 0.74 | 4.86 ± 0.30 | 86.97 ± 5.66 | 2.37 ± 0.37 |
| Hatched | FRD | Feather River | HZ | 0.22 ± 0.16 | 9.18 ± 3.08 | 0.71 ± 0.10 | 1076.67 ± 15.28 | 10.57 ± 1.28 |
| Hatched | BCC | Battle&Butte | HZ | 0.13 ± 0.02 | 3.31 ± 0.91 | 0.28 ± 0.17 | 187.33 ± 6.43 | 2.55 ± 0.22 |
| Hatched | SAC | Sacramento River | HZ | | 44.37 ± 18.54 | 4.98 ± 0.88 | 2623.33 ± 95.04 | 32.63 ± 4.27 |
| Hatched | SSC | Sacramento River | HZ | | 10.12 ± 2.23 | | 973.00 ± 74.18 | 13.83 ± 1.22 |
| Hatched | NFB | Battle&Butte | HZ | | 23.47 ± 10.09 | | 1556.67 ± 105.99 | 46.23 ± 6.67 |
| Hatched | SFB | Battle&Butte | HZ | 0.19 ± 0.07 | 6.36 ± 1.19 | | 657.67 ± 9.29 | 9.36 ± 0.76 |
| Hatched | CCH | Clear Creek | HZ | 0.21 ± 0.14 | 2.85 ± 1.38 | 0.67 ± 0.20 | 194.67 ± 19.63 | 2.87 ± 0.27 |
| Hatched | CCM | Clear Creek | HZ | | 16.97 ± 5.14 | | 465.33 ± 19.43 | 17.40 ± 1.32 |
| Pre-Spawn | FRU | Feather River | SW | 0.15 ± 0.02 | 1.87 ± 0.97 | 11.83 ± 0.71 | 29.40 ± 3.56 | |
| Pre-Spawn | FRD | Feather River | SW | 0.45 ± 0.10 | 3.36 ± 0.23 | 8.25 ± 0.25 | 96.53 ± 7.34 | 1.26 ± 0.21 |
| Pre-Spawn | BCC | Battle&Butte | SW | 0.39 ± 0.11 | 2.11 ± 0.16 | 4.20 ± 0.26 | 371.67 ± 15.50 | |
| Pre-Spawn | SAC | Sacramento River | SW | 0.08 ± 0.03 | 1.23 ± 0.09 | 2.91 ± 0.02 | 70.57 ± 4.68 | |
| Pre-Spawn | SSC | Sacramento River | SW | 0.11 ± 0.05 | 1.86 ± 0.03 | 8.25 ± 0.21 | 37.97 ± 1.66 | |
| Pre-Spawn | NFB | Battle&Butte | SW | 0.15 ± 0.05 | | 0.67 ± 0.03 | 259.67 ± 10.07 | |
| Pre-Spawn | SFB | Battle&Butte | SW | 0.23 ± 0.06 | 2.77 ± 0.16 | 3.05 ± 0.26 | 88.50 ± 0.95 | 2.21 ± 0.13 |
| Pre-Spawn | CCH | Clear Creek | SW | 0.21 ± 0.08 | 1.90 ± 0.10 | 7.76 ± 0.39 | 55.13 ± 6.07 | |
| Pre-Spawn | CCM | Clear Creek | SW | 0.23 ± 0.07 | 1.23 ± 0.36 | 9.31 ± 0.53 | 29.13 ± 2.42 | |
| Spawn | FRU | Feather River | SW | 0.72 ± 0.05 | 7.54 ± 1.00 | 31.73 ± 0.93 | 103.27 ± 15.67 | 10.84 ± 1.61 |
| Spawn | FRD | Feather River | SW | 0.52 ± 0.02 | 5.38 ± 0.30 | 38.23 ± 0.71 | 31.07 ± 4.10 | 4.72 ± 0.21 |
| Spawn | BCC | Battle&Butte | SW | 0.30 ± 0.04 | 2.56 ± 0.51 | 10.39 ± 0.81 | 37.83 ± 7.00 | |
| Spawn | SAC | Sacramento River | SW | 0.21 ± 0.03 | 1.62 ± 0.23 | 2.75 ± 0.43 | 237.33 ± 26.01 | |
| Spawn | SSC | Sacramento River | SW | 0.22 ± 0.04 | 1.76 ± 0.67 | 27.80 ± 0.35 | 272.67 ± 16.29 | |
| Spawn | NFB | Battle&Butte | SW | 0.28 ± 0.09 | 4.47 ± 0.75 | 12.03 ± 0.42 | 49.63 ± 2.66 | |
| Spawn | SFB | Battle&Butte | SW | 22.30 ± 12.51 | 418.33 ± 49.89 | 1333.33 ± 102.63 | 3436.67 ± 94.52 | 338.33 ± 40.81 |
| Spawn | CCH | Clear Creek | SW | 0.41 ± 0.09 | 1.60 ± 0.72 | 3.48 ± 0.35 | 119.00 ± 11.53 | |
| Spawn | CCM | Clear Creek | SW | 0.33 ± 0.11 | 1.09 ± 0.27 | 12.50 ± 1.13 | 16.67 ± 2.38 | |
| Incubation | FRU | Feather River | SW | 0.20 ± 0.09 | 3.26 ± 0.12 | 6.77 ± 0.20 | 932.33 ± 49.33 | |
| Incubation | FRD | Feather River | SW | 0.28 ± 0.13 | 2.26 ± 0.53 | 8.98 ± 0.47 | 166.67 ± 20.82 | |
| Incubation | FR2 | Feather River | SW | 0.33 ± 0.05 | 1.77 ± 0.10 | 11.10 ± 0.50 | 140.33 ± 9.29 | |
| Incubation | FR3 | Feather River | SW | 0.23 ± 0.12 | 1.21 ± 0.19 | 12.77 ± 0.61 | 23.57 ± 1.55 | |
| Incubation | BCC | Battle&Butte | SW | 0.11 ± 0.04 | 0.52 ± 0.04 | 4.16 ± 0.12 | 156.33 ± 4.73 | |
| Incubation | SAC | Sacramento River | SW | 0.17 ± 0.01 | 2.91 ± 0.07 | 12.53 ± 0.70 | 258.67 ± 30.86 | |
| Incubation | SSC | Sacramento River | SW | 0.19 ± 0.04 | 2.48 ± 0.60 | 14.80 ± 0.75 | 44.47 ± 4.57 | |
| Incubation | NFB | Battle&Butte | SW | 0.23 ± 0.07 | 1.39 ± 0.06 | 7.60 ± 0.37 | 166.33 ± 5.69 | |
| Incubation | SFB | Battle&Butte | SW | 0.33 ± 0.12 | 2.81 ± 0.53 | 7.25 ± 0.44 | 456.00 ± 23.07 | 3.14 ± 0.48 |
| Incubation | CCH | Clear Creek | SW | 0.36 ± 0.09 | 2.94 ± 0.16 | 14.37 ± 0.12 | 426.67 ± 18.15 | |
| Incubation | CCM | Clear Creek | SW | 0.47 ± 0.10 | 0.96 ± 0.19 | 7.70 ± 0.42 | 952.67 ± 59.41 | |
| Hatched | FRU | Feather River | SW | 0.30 ± 0.15 | 2.20 ± 0.16 | 11.40 ± 1.08 | 33.30 ± 6.10 | |
| Hatched | FRD | Feather River | SW | 0.68 ± 0.07 | 1.63 ± 0.97 | 6.45 ± 0.38 | 31.87 ± 3.01 | |
| Hatched | BCC | Battle&Butte | SW | 0.07 ± 0.02 | 0.45 ± 0.24 | 0.61 ± 0.02 | 52.63 ± 0.50 | |
| Hatched | SAC | Sacramento River | SW | 0.09 ± 0.02 | 0.96 ± 0.17 | 1.63 ± 0.15 | 129.33 ± 12.06 | |
| Hatched | SSC | Sacramento River | SW | 0.09 ± 0.02 | 0.66 ± 0.10 | 13.60 ± 0.52 | 49.57 ± 2.25 | |
| Hatched | NFB | Battle&Butte | SW | 0.06 ± 0.02 | 0.87 ± 0.12 | 2.98 ± 0.20 | 38.97 ± 3.20 | |
| Hatched | SFB | Battle&Butte | SW | 0.22 ± 0.04 | 0.84 ± 0.37 | 3.84 ± 0.32 | 22.70 ± 4.00 | 1.02 ± 0.41 |
| Hatched | CCH | Clear Creek | SW | 0.36 ± 0.08 | 1.61 ± 0.62 | 8.17 ± 0.70 | 54.97 ± 4.97 | |
| Hatched | CCM | Clear Creek | SW | 0.28 ± 0.07 | 0.70 ± 0.52 | 9.70 ± 1.25 | 44.13 ± 7.12 | |
